# First passage time analysis of spatial mutation patterns reveals evolutionary dynamics of pre-existing resistance in colorectal cancer

**DOI:** 10.1101/2022.02.22.481463

**Authors:** Magnus J. Haughey, Aleix Bassolas, Sandro Sousa, Ann-Marie Baker, Trevor A. Graham, Vincenzo Nicosia, Weini Huang

## Abstract

The footprint left by early cancer dynamics on the spatial arrangement of tumour cells is poorly understood, and yet could encode information about how therapy resistant sub-clones grew within the expanding tumour. Novel methods of quantifying spatial tumour data at the cellular scale are required to link evolutionary dynamics to the resulting spatial architecture of the tumour. Here, we propose a framework using first passage times of random walks to quantify the complex spatial patterns of tumour cell population mixing. First, using a toy model of cell mixing we demonstrate how first passage time statistics can distinguish between different pattern structures. We then apply our method to simulated patterns of wild-type and mutated tumour cell population mixing, generated using an agent-based model of expanding tumours, to explore how first passage times reflect mutant cell replicative advantage, time of emergence and strength of cell pushing. Finally, we analyse experimentally measured patterns of genetic point mutations in human colorectal cancer, and estimate parameters of early sub-clonal dynamics using our spatial computational model. We uncover a wide range of mutant cell replicative advantages and timings, with the majority of sampled tumours consistent with boundary driven growth or short-range cell pushing. By analysing multiple sub-sampled regions in a small number of samples, we explore how the distribution of inferred dynamics could inform about the initial mutational event. Our results demonstrate the efficacy of first passage time analysis as a new methodology for quantifying cell mixing patterns *in vivo*, and suggest that patterns of sub-clonal mixing can provide insights into early cancer dynamics.

## Introduction

Understanding the origins and effects of intra-tumour heterogeneity is a fundamental challenge in cancer research and is critical for managing therapy resistance and improving treatment outcomes [1–5]. Detectable tumours represent a diverse population of cells, often containing sub-populations, or *sub-clones*, carrying genetic alterations which may confer resistance to targeted cancer therapy. Anti-cancer treatment alters the selective landscape within the tumour, enabling pre-existing resistant sub-clones to expand, which unavoidably leads to relapse and treatment failure [2, 6]. Understanding the evolution of resistant sub-clones within tumours will therefore greatly advance our ability to mitigate treatment resistance and even exploit evolutionary principles to improve the efficacy of cancer treatment [7].

Direct observation of the early evolution of resistance is often infeasible as, in some cancers, sub-clones arise in the early, undetectable, malignancy [4, 8]. This problem is confounded by the difficulty of obtaining temporal samples in humans. Recent computational and mathematical modelling, however, has been applied to estimate the probability of developing treatment-resistant sub-clones, and the influence of tissue architecture on the evolution of both neutral and oncogenic mutations [6, 9–11]. Spatial information itself is used in cancer diagnosis and prognosis [12–14]; for example, the spatial distribution of tumour infiltrating immune cells has been shown to have prognostic value in some cancers [15–19]. In this study, we reason that the sub-clonal spatial pattern is a readout of the evolutionary history of a tumour and could present a route to quantifying the evolution of treatment resistance. To investigate this, we use spatial computational modelling to develop methodologies for inferring the growth history of tumour sub-clones based on their spatial arrangement in sampled tumours.

We apply our analysis to human colorectal cancer samples, where the spatial composition of point mutations driving treatment resistance at the cellular level are mapped by the BaseScope RNA *in situ* hybridisation assay. Previously, Baker *et al*. used this method to reveal diverse patterns of sub-clonal mixing between mutated (resistant) and wild-type tumour cells (Fig 1a), and quantified these patterns using a spatial analogue of Shannon’s entropy [20]. Initial spatial analysis suggested that the observed BaseScope patterns were consistent with early arising, weakly selective, mutated sub-clones or later arising sub-clones endowed with a larger replicative advantage over the original tumour (wild-type) population. Whilst Shannon’s entropy provided the first analysis of tumour heterogeneity, it is defined at a single spatial scale and thus is strongly scale dependent. We address this issue here by quantifying clustering and heterogeneity of cell mixing patterns at multiple spatial scales using the statistics of random walks to obtain a more complete description of the complex sub-clonal patterns elucidated with BaseScope.

**Fig 1.**
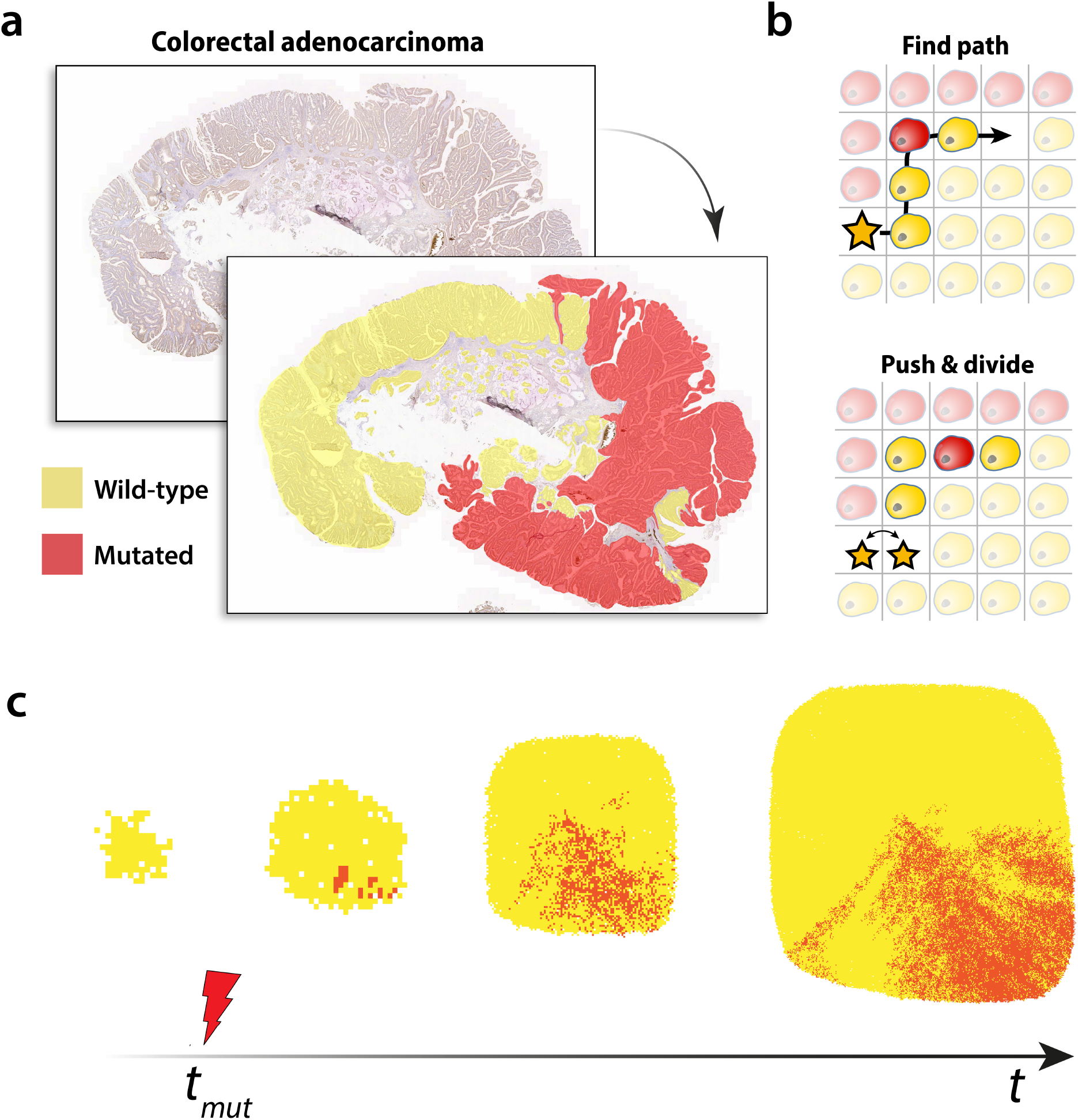
Sub-clonal mixing in human colorectal tumours. **(a)** Sub-clonal mixing patterns in human colorectal adenocarcinoma tissue revealed with BaseScope. Tumour cells carrying the *KRAS* G12A mutation are highlighted in red. Those which are wild-type at this loci are highlighted in yellow. **(b)** Cell pushing in spatial simulations. A dividing cell (star) creates the space needed to divide into two cells by pushing neighbouring cells along a path to a nearby empty lattice point. **(c)** Computational model of tumour sub-clonal mixing. Competing wild-type (yellow) and mutated (red) cell populations clonally expand on a 2-dimensional square lattice.

Many spatial systems can be naturally represented as a network of interacting and connected nodes, of different classes, as a way to study the effects of the system’s structure on its dynamics. Heterogeneity and segregation within networks can influence the statistics of a random walker embedded in the graph, making random walks a useful tool in network science to quantify the structural properties of a system [21–24]. By quantifying of the structure of the sub-clonal patterns at multiple length scales, therefore improving on previous analysis of these patterns, we propose that methods exploiting random walks have the potential to advance our understanding of the link between spatial arrangement of sub-clones and the underlying sub-clonal dynamics.

In this study, we focus on the class mean first passage times (CMFPT) on a network, defined as the expected time, *τ*_*αβ*_, for a random walker beginning on a node of class *α* to first arrive at a node of class *β*. This method has recently been applied to other complex systems to quantify spatial segregation in voting patterns [25], ethnic segregation in UK and US metropolitan areas [26], and to show that internal clustering and spatial heterogeneity of individuals of different ethnic groups contributed to the observed excess of infectious diseases [27]. Here, we leverage the CMFPT to quantify the complex spatial patterns of mutated sub-clonal cells in BaseScope images.

We first demonstrate the capability of the CMFPT method to measure pattern structures using an artificial toy model of wild-type (WT) and mutant population mixing. We then compare CMFPT measurements of sub-clonal mixing patterns in human colorectal cancers to the same measurements derived from our agent-based simulations. We perform parameter estimation using our computational model, predicting the relative age and replicative fitness advantage of the mutated sub-clone and the strength of cell pushing most consistent with the spatial patterns observed with BaseScope. Our analysis indicates that the resistant sub-clones often appear relatively early in the expansion of the WT population, and exhibit a wide range of fitness advantages over WT cells. This work demonstrates the capability of the class mean first passage time as a method of quantifying cell mixing patterns *in vivo*, and our findings suggest that patterns of sub-clonal mixing in mature tumours can provide insights into early sub-clonal dynamics.

## Results

### Computing first passage time statistics

To estimate the class mean first passage time (CMFPT), we simulate the trajectory of random walkers on cell mixing patterns embedded in a square lattice. The mean number of steps, *τ*_*αβ*_, taken to first arrive at a node of class *β* when departing from a node of class *α*, is computed over all starting nodes of the same class *α* on the graph. In this study we focus on spatial patterns of two classes, mutated (red) and wild-type cells (yellow), giving rise to four possible CMFPT quantities: *τ*_*rr*_, *τ*_*ry*_, *τ*_*yr*_ and *τ*_*yy*_. Different spatial patterns will differ naturally in magnitude and shape. Thus, to compare the CMFPT of different images we normalise the first passage times for each image to the corresponding quantities derived under a null model, in which the constituent classes in the spatial pattern are reassigned uniformly at random to each lattice point, but maintaining the abundances of each class and the general topology of the original image. We denote these quantities as the normalised first passage times 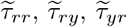 and 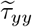.

We explore the phase space spanned by 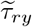 and 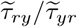, where 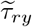 and 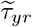 denote the normalised CMFPT for red→yellow and yellow→red transitions respectively. Through our analysis we find that the combination of these quantities enables the segmentation of spatial patterns which differ in heterogeneity, and separates images containing clusters of different characteristic sizes. Measurements in this phase space have been used in previous analyses involving CMFPT to characterise colour distributions on 2-dimensional lattices [25]. Spatial patterns in which the two classes are distributed uniformly at random will lie at 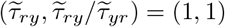, since this particular pattern exactly resembles the null model used for normalisation. In general, patterns which contain a more ordered, or segregated, distribution of classes will give rise to 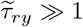. Those which contain large clusters, or percolating structures, of yellow classes will appear in the region 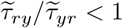 and, conversely, those which contain similar structures of red classes will appear in the region 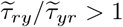.

We define the class ratio, *ϕ*, of an image as the ratio of red to yellow classes,

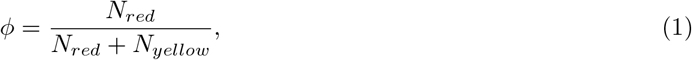

where *N*_*red*_ and *N*_*yellow*_ represent the number of red and yellow pixels in an image respectively. Note while *ϕ* is an direct input in our toy model, it is an outcome of the agent-based simulations, depending on the replicative advantage of the sub-clonal population, the relative time at which it emerges in the simulations, and the strength of cell pushing.

### A toy model of 2-dimensional cell mixing patterns

We first develop a toy model to generate a set of diverse spatial patterns of two colours (red and yellow) in a 2-dimensional lattice background. In particular, we generate three distinct patterns, which are: (1) small clusters; (2) large centred cluster and (3) column arrangement (Fig 2a-f). We run a number of simulations whilst varying class ratio, *ϕ*, between 0 ≤ *ϕ* ≤ 1 for each of these patterns and calculate the corresponding normalised CMFPT in the 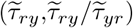 phase space in each case (Fig 2g). Results from these initial calculations demonstrate how pattern structure and class ratio are reflected in the CMFPT, with a clear correspondence between *ϕ* and 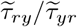 value (vertical axis) observed across all patterns. Patterns with a 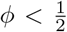 generally lie below 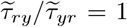, and those with a 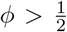 fall above 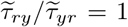, reflecting the increasing segregation of classes at either extreme of class ratio. Moreover, this effect is intensified for specific spatial patterns, with stronger dependence for fully segregated patterns (*i.e*. column patterns *e, f* in Fig 2g), and a weaker dependence for more intermixed patterns (*i.e*. clusterised patterns *a, b* in Fig. 2g).

**Fig 2.**
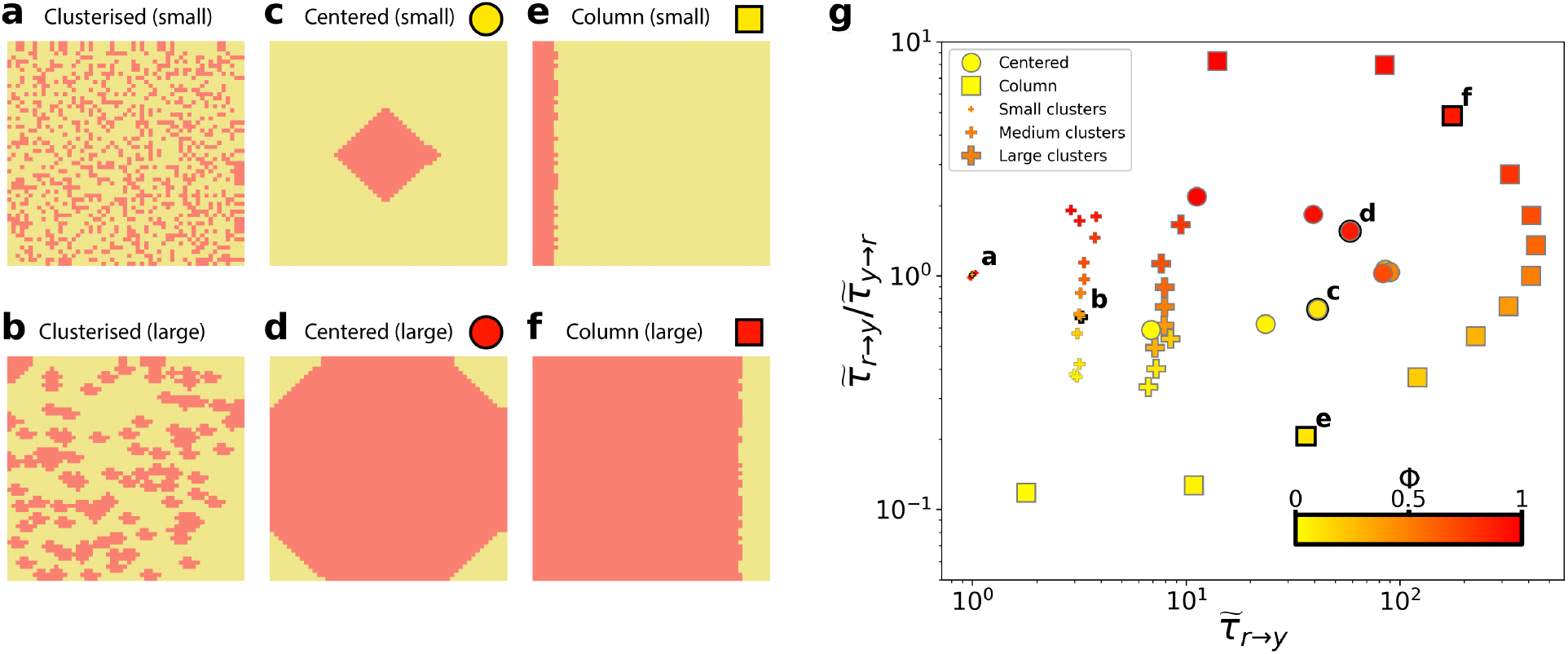
Characterising 2-dimensional toy model mixing patterns using CMFPT. Patterns were generated using the (**a, b**) clusterised; (**c, d**) centred and (**e, f**) column models. **(g)** Location of the patterns obtained for the three models with varying class ratio, *ϕ*.

Our normalised CMFPT measure allows for more precise interrogation of pattern structures than class ratio alone. The separation of each pattern type is shown along the 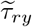 (horizontal) direction of Fig 2g. Being most similar to its corresponding null model used to normalise the CMFPT, the small clusterised pattern (Fig 2a) occupies a region of the phase space near to 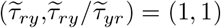. The normalised CMFPT measurement is sensitive to an increase in the cluster size, even when class ratio remains constant (Fig 2b). For both the centred (Fig 2c, d) and column (Fig 2e, f) patterns, discrete trajectories in the phase space are observed in Fig 2g, with the value of *ϕ* determining the position of the specific pattern along the curve.

### Analysing simulated tumour sub-clone mixing patterns

We next investigate the efficacy of our method in characterising cell mixing patterns generated *in silico*. We developed a 2-dimensional agent-based simulation model of expanding tumour populations, incorporating cell birth, death and mutation on a lattice (Fig 1b, c). Simulations begin with a single wild-type (WT) tumour cell in the centre of the lattice which seeds a growing WT population. After a threshold number of WT cell divisions, specified by the model parameter *t*_*mut*_, an existing WT cell acquires a mutation which confers a fitness advantage of *s* (see Methods). We model fitness via a modulation of cell replication rates, such that cells carrying the mutation have a replication rate which is (*s* × 100)% greater than that of WT cells. Neutral selection, where the mutant sub-clone grows at the same rate as the WT population, corresponds to *s* = 0. These simulations result in 2-dimensional spatial patterns of WT cells coexisting with a sub-clonal population of mutated cells. We utilise this framework to explore the impact of varying three model parameters, i.e. mutant replicative advantage, *s*, the relative time, *t*_*mut*_, at which the mutation appears in the growing WT population, and the strength of cell-cell pushing on the lattice, *q* (see Table S1).

Before analysing the patterns of the normalised CMFPT as a function of *s, t*_*mut*_ and *q*, we first characterise its relationship with the class ratio, *ϕ*, in each of the patterns (Fig 3). Similar to our observation in the toy model, we find a complex non-linear relationship between *ϕ* and the CMFPT, and a general trend of increasing 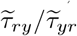 with class ratio. In the toy model, we can vary *ϕ* and the underlying pattern independently, since *ϕ* is an explicit model parameter. In our agent-based tumour model, however, this ratio has a dependence on *s, t*_*mut*_ and *q*, being simply a readout quantity rather than a tunable parameter. Thus, class ratio and pattern structure are naturally coupled. Correspondingly, certain underlying pattern structures are observed only for small mutant sub-clones, and therefore small *ϕ* (Fig 3d), whereas combinations of (*s, t*_*mut*_, *q*) which give rise to higher *ϕ* will also result in substantially different pattern topologies (Fig 3a). As a result, the CMFPT measurements of the simulated patterns do not fall onto distinct curves (Fig 3e), as was observed for different underlying structures in the toy model.

**Fig 3.**
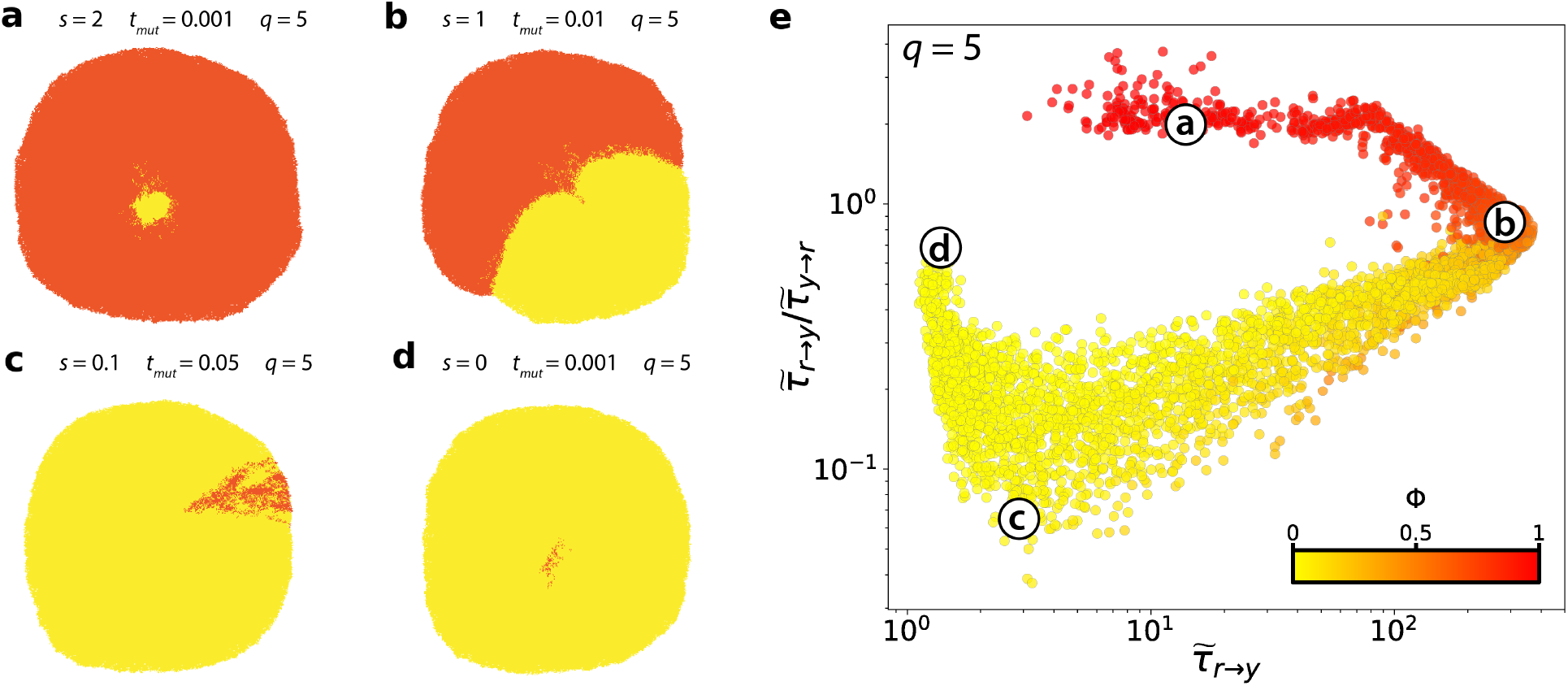
Analysis of simulated sub-clonal mixing patterns. **(a-d)** Simulated sub-clonal mixing patterns with cell pushing strength of *q* = 5. **(e)** Analysis of all simulated mixing patterns simulated with a cell pushing value of *q* = 5 in the 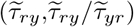 phase space. Point colour corresponds to the relative abundance of red and yellow, *ϕ*.

Beyond class ratio, we investigate the extent to which the CMFPT reflects spatial patterns generated under different model parameter combinations (Fig 4). When we classify our simulations according to mutant fitness, *s*, and the arising time of the mutant, *t*_*mut*_, spatial patterns generated by different parameter combinations occupy distinct regions of the 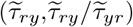 phase space, demonstrating that the CMFPT can be used to quantify distinct spatial heterogeneity. Patterns corresponding to neutral selection, *s* = 0 (blue points in Fig 4), lie in the leftmost region 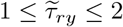, and sub-clones with strong selection, 2 ≤ *s* ≤ 3 (yellow and red points in Fig 4), are clearly segregated at larger values of 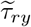. Due to the coupled effects of *s* and *t*_*mut*_ on the final class ratio, tumours with early arising, weakly selected mutant sub-clones (Fig 4, small blue points), and those with later arising, strongly selected sub-clones (Fig 4, large red points) share similar values of *ϕ* (shown by colours in Fig 4a-d, inset). Interestingly, despite having a similar mutant frequency, *ϕ*, the CMFPT is sensitive to the topological differences between patterns generated in these two contrasting scenarios. Tumours with the earliest arising sub-clones occupy a unique area of the phase space for all cell pushing strengths.

**Fig 4.**
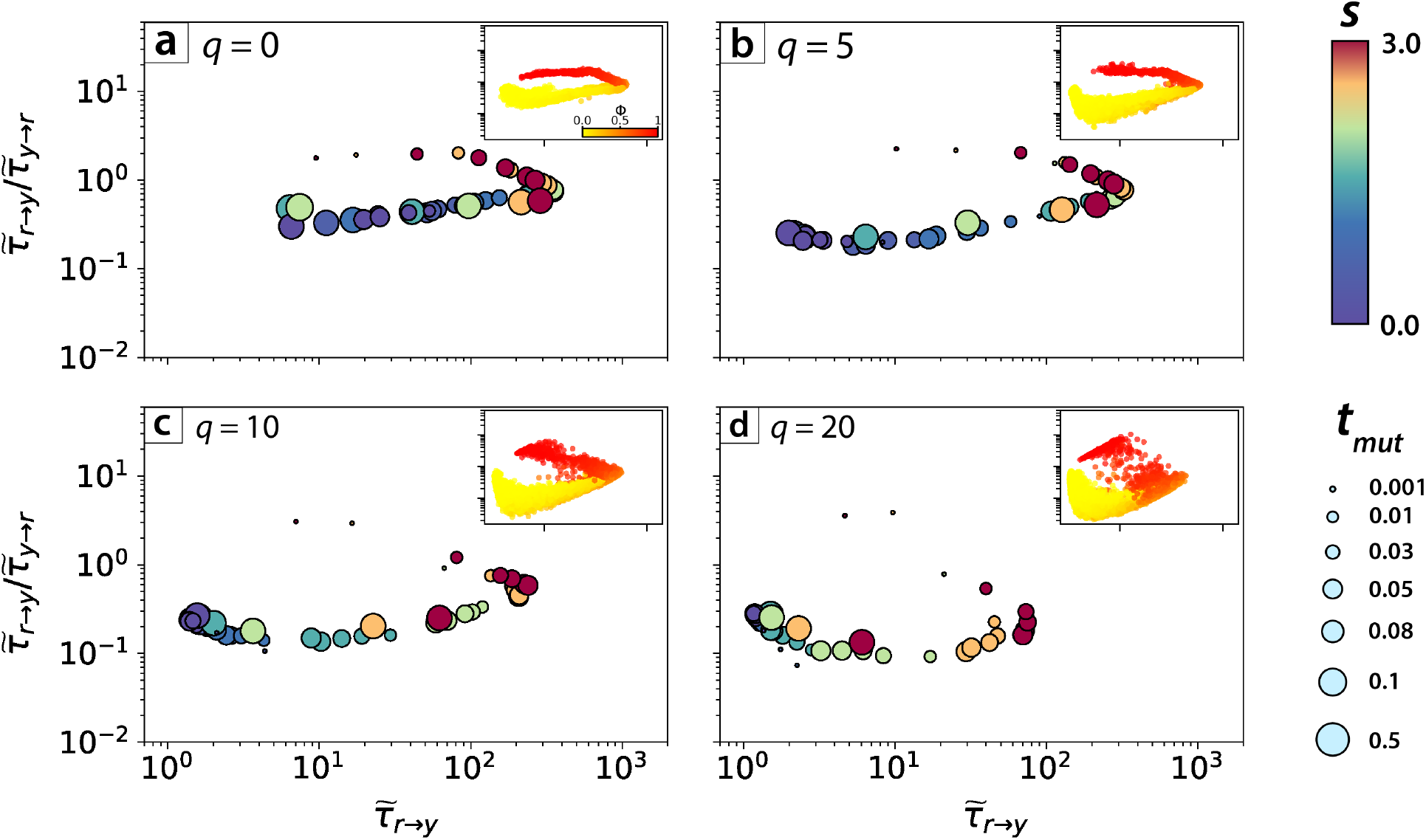
Analysis of simulated sub-clonal mixing patterns for varying sub-clonal dynamics. Mean CMFPT values for simulated sub-clonal mixing patterns in the phase space 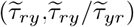, with points coloured according to values of model parameter *s* and point size depending on parameter *t*_*mut*_. Images are separated depending on their pushing value *q* = 0 (**a**), *q* = 5 (**b**), *q* = 10 (**c**) and *q* = 20 (**d**).

Varying the strength of cell pushing, *q*, has clear qualitative impacts on spatial heterogeneity. At weak pushing strengths cell displacement occurs at short ranges, leading to greater clustering of sub-clones. Conversely, at larger values of *q* cells are subject to more frequent displacement by nearby dividing cells, so that any sub-clonal clusters are more quickly dispersed. These effects are also reflected in the distribution of CMFPT measurements in the 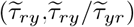 phase space (Fig 4a-d). For weak pushing strengths (Fig 4a) the simulated spatial patterns follow a path in phase space more similar to that of the centred toy model patterns (Fig 2c, d), with two distinct “arms” demarcating patterns with *ϕ <* 0.5 and those with *ϕ >* 0.5: the intersection of these arms reflecting the theoretical upper limit of 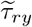 when both mutant and WT populations arranged in a fully separated, semi-circular clusters with *ϕ* = 0.5. In the limit of zero cell pushing (*q* = 0), varying the mutant replicative advantage, *s*, and arising time, *t*_*mut*_, impacts the final class ratio. Since, however, cell proliferation is limited to the periphery of the system, variations in these parameters have little influence on the spatial heterogeneity of the resulting sub-clonal patterns. This lack of variation in sub-clonal pattern topology at weaker *q* is reflected through a low variance in the measured CMFPT as *s* and *t*_*mut*_ are varied, and measured values cluster strongly in the phase space according to the class ratio of the sub-clonal pattern (Fig 4a, inset).

On the contrary, under stronger cell pushing the diversity in the observed sub-clonal pattern structures is greater, resulting in increased variance in the distribution of CMFPT values and weaker clustering of CMFPT measurements according to the class ratio of the simulated pattern. This is explained by the loss of distinct clusters of closely related cells when cell pushing strength is increased. The increased fragmentation of subclones is captured by the CMFPT and represented by a steeper initial decline in 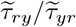 for patterns with low *s*.

### Quantifying sub-clonal dynamics in human colorectal cancer

Having characterised the performance of the CMFPT measure with both toy model and agent-based simulated tumours, we now apply our method to human colorectal carcinoma samples spatially mapped by BaseScope [20]. We analysed 22 cell mixing patterns produced with BaseScope (Fig S1a) which detail the spatial composition of tumour cells, mapping out sub-clonal populations carrying point mutations in one of three genes commonly mutated in colorectal cancer – *PIK3CA, BRAF* and *KRAS*. Mutations in these genes have been associated with cancer driver events, poor prognosis and treatment resistance [28–32]. Based on our understanding of how the CMFPT measurement relates to sub-clonal mixing patterns generated with our agent-based model, we aim to elucidate the underlying dynamics driving the early growth of the mutated sub-clones observed with BaseScope. It is important to note that, whilst the mutation time in our 2-dimensional simulations, *t*_*mut*_, describes the time at which the mutation first appears in the entire system, the interpretation of this parameter changes when applied to the BaseScope samples. It is extremely rare that the same mutation will happen independently in multiple regions of any tumour, yet since the BaseScope images represent 2-dimensional sub-samples of a larger 3-dimensional system, spatially distant sub-clonal regions are often observed. Accordingly, any inferred values of *t*_*mut*_ will represent the relative length of time that the WT population was expanding in that localised area before the invasion of the mutant sub-clone. In a few cases, where the BaseScope samples involve multiple spatially discontinuous tumour areas, the distribution of inferred *t*_*mut*_ values of all local areas reveals how fast the mutant population spread in these tumours (details explained below).

Initial analyses of the BaseScope images (Fig S7), for which we estimated the CMFPT over the totality of each image, highlighted some important additional processing steps that should be applied to the BaseScope images prior to parameter inference. Crucially, the presence of un-highlighted areas in the BaseScope images, representing all other non-cancerous tissue, significantly impacted our estimations of the CMFPT. We do not consider non-cancerous cells in our agent-based model, to maintain modest model complexity, making comparisons to simulated sub-clonal patterns uninformative. To improve comparisons between experimental and simulated sub-clonal patterns, we execute a number of pre-processing steps on the BaseScope images. We first process each image by manually filling in un-highlighted interior area of colonic crypts in predominantly WT or mutant regions with the relevant colour (Fig S1b). To address the remaining un-highlighted areas which correspond to regions of non-cancerous cells (*e.g*. connective tissue), we separate those BaseScope images with spatially discontinuous tumour regions into smaller sub-sections, where each is made of contiguous regions of WT and mutated cells. By analysing these sections individually, we reduce the influence of un-highlighted space separating these regions, enabling a better comparison of experimental and simulated measurements.

Following these pre-processing steps, the results are largely improved compared to our initial analyses of the full BaseScope images, and reveal a similar general trend to that observed from our *in silico* pattern results (Fig 5). With the implementation of additional pre-processing steps, we observe a good correspondence between experimental and model data in terms of *ϕ*, with higher *ϕ* patterns appearing in the 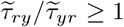 region of the phase space, and lower *ϕ* patterns situated below 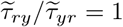. Only a handful of the analysed BaseScope patterns lie below 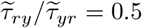, a region of the phase space predominantly occupied by simulated patterns with high cell pushing, *q*, and low mutant selection, *s*, suggesting cell pushing is generally weak *in vivo*.

**Fig 5.**
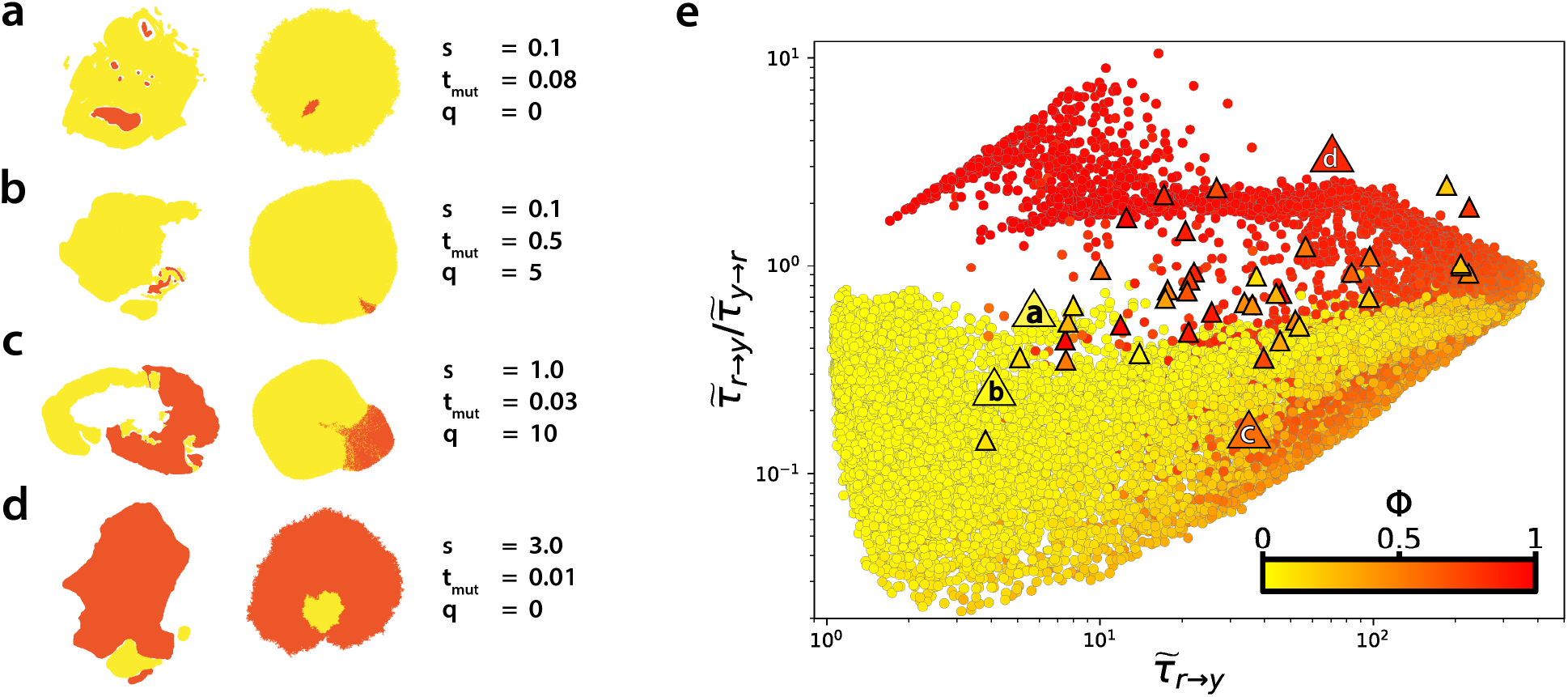
Analysing sub-components of BaseScope sub-clonal patterns. **(a-d)** Representative examples of sub-components of the BaseScope samples (after pre-processing steps applied) and their best-fit simulated sub-clonal patterns. **(e)** CMFPT analysis of all simulated sub-clonal patterns (circles) and connected WT and mutant cell sub-regions within BaseScope images (triangles). Points are coloured according to pattern class ratio, *ϕ*.

**Fig 6.**
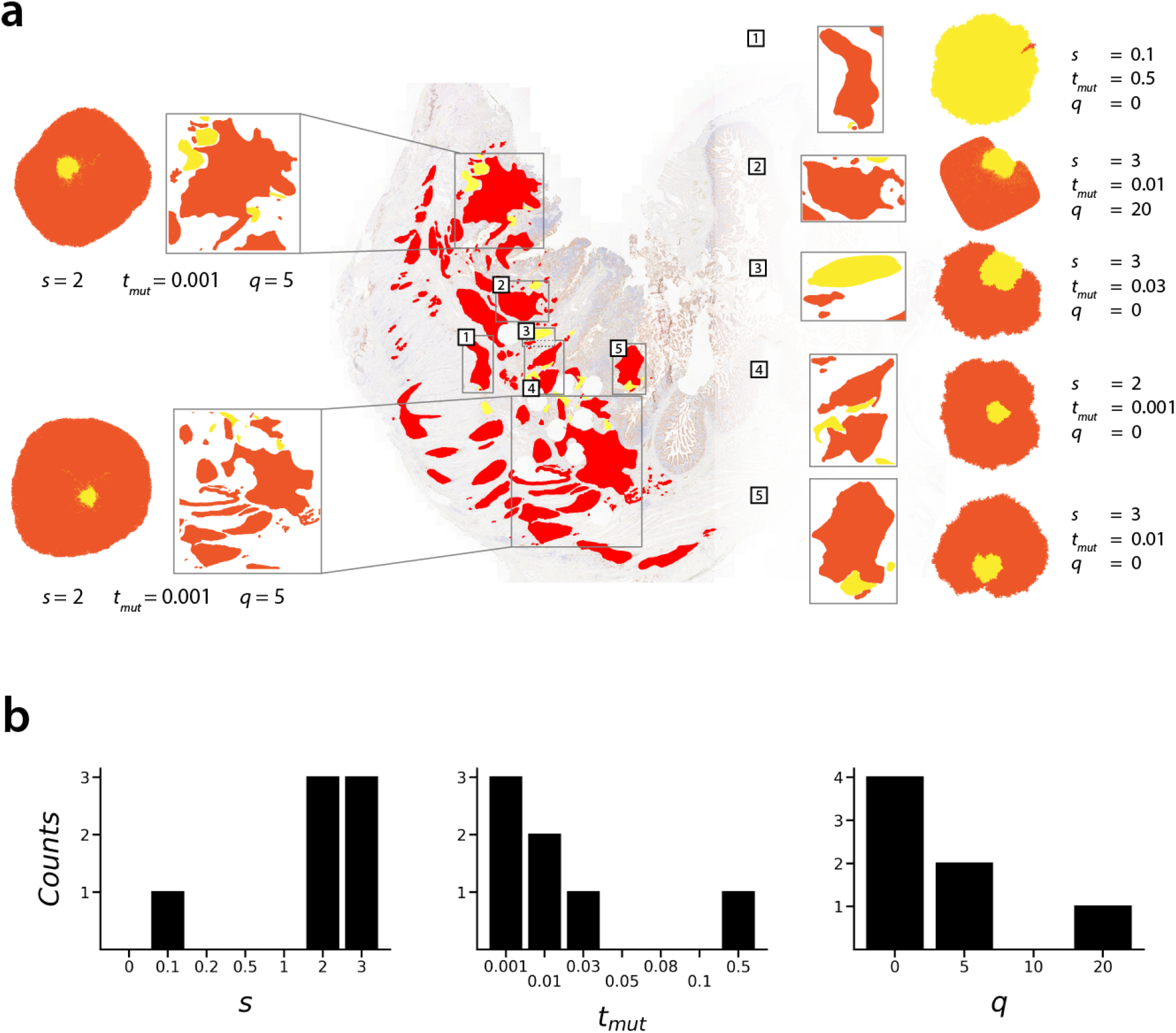
**(a)** Sub-section analysis of BaseScope sample 34. Best-fit simulated sub-clonal pattern shown next to each sub-section with corresponding model parameters. **(b)** Marginal distributions of median inferred model parameters.

To describe the sub-clonal dynamics more quantitatively, we performed a grid-search to find the best-fit simulated parameters for each of the analysed sub-components in the BaseScope images (Figs 5a-d, Figs S8-S23) (see Methods). Plotting the best-fit simulated patterns demonstrates that our method not only captures the statistical features of the experimental images, but that the best-fit simulated patterns often also share a close visual correspondence to the BaseScope patterns both in terms of class ratio and pattern heterogeneity. BaseScope patterns with high segregation of WT and mutant populations are predicted to have low cell pushing. Highly segregated patterns which, in addition, have a large mutant frequency, *ϕ*, are most consistent with model patterns simulated with early arising, mutant sub-clones endowed with a strong replicative advantage.

The inferred best-fit parameters for connected WT and mutant regions within our BaseScope images point to a wide range of sub-clonal growth dynamics. Analysis of the majority of the BaseScope samples suggests that the mutated populations experience some degree of replicative advantage, *s*, over the WT population, with some patterns consistent with simulated tumours generated in the parameter range 1 ≤ *s* ≤ 3. Inferred values for the relative time of mutant sub-clone invasion, *t*_*mut*_, within each BaseScope sub-component range between 0.1% and 10% for the majority of the samples, suggesting that the mutant sub-clones arise early in the expansion of the WT tumour populations. Consistent with our earlier supposition, the inferred value of cell pushing parameter *q* is mostly *q* = 0 or *q* = 5, however some larger inferred pushing values are predicted.

For three of the most fragmented BaseScope images, samples 02, 28 and 34, we were able to separately analyse several isolated sub-components of tumour cells and construct a distribution of inferred parameter values across the full images (Figs S32, S33 & 6 respectively). We surmise that the distribution of inferred selection strength, *s*, and mutation time, *t*_*mut*_, can offer additional insights into the nature of the early sub-clonal evolution in these tumours. As previously mentioned, *t*_*mut*_ should not be interpreted as the time at which the mutant population first appears in the tumour, but instead it indicates the relative length of time that the WT population was expanding in that area prior to invasion by the expanding mutated sub-clone. Despite this, there will be some correspondence between the distribution of inferred local *t*_*mut*_ values and the time of the original mutational event: with our computational modelling demonstrating that earlier arising mutations tend to lead to higher infiltration of the mutant sub-clone throughout the mature tumour, and less variegation in the mutant sub-clone pattern. Following this rationale, we would expect an earlier mutational event to result in a unimodal distribution of inferred *t*_*mut*_ values, with a low variance. Conversely, later occurring mutations should lead to a wider distribution in inferred *t*_*mut*_ values, which may deviate from unimodality.

Samples 02 and 34 (Figs S32 and 6 respectively) reveal a trend of high mutant replicative advantage and early emerging sub-clones. Both possess a narrow distribution of inferred *t*_*mut*_ across the different sub-sampled regions, concentrated towards low *t*_*mut*_ values, indicating early invasion of the mutant sub-clone within the WT population, and pointing to an overall early emerging mutation in the evolution of the tumour. In both cases, the narrow distribution of *t*_*mut*_ is coupled to a narrow distribution of inferred selection strength, *s*, trending towards large values. These factors point to early arising, highly positively selected mutant sub-clones within these tumours. Contrasting these two samples, BaseScope sample 28 (Fig S33) contains a larger proportion of sub-regions consistent with a late invading mutant sub-clone (*t*_*mut*_ ≥ 0.1), and a wider distribution of inferred *t*_*mut*_ values, along with lower inferred selection strength. Whilst we have limited resolution in these distributions, this might suggest that the mutation itself occurred later in the evolution of sample 28 than it did in samples 02 and 34.

The majority of the analysed sub-regions within BaseScope samples 02, 28 and 34 suggest an overall trend of weak cell pushing, *q*, with many inferred pushing values of *q* = 0. This trend persists across all analysed BaseScope samples, with zero cell pushing (*q* = 0) best describing over 50% of all patterns and *q* = 0 or *q* = 5 accounting for the majority of inferred pushing strengths. It should be noted, however, that inference of this parameter may be affected most by the sub-sampling of sub-clonal patterns, which might bias inference to lower pushing values.

The fact that BaseScope samples 02 and 34 in particular point to similar sub-clonal dynamics may not be coincidental. The high degree of tumour cell fragmentation and mixing within non-cancerous tissue meant that it was necessary to sample a number of smaller sub-regions within these images, in order to achieve better concordance between experimental and simulated data. It could be that the dynamics of early emerging, highly replicative mutant sub-clones was responsible for producing the fragmented tumour cell population which was observed *in vivo*. In fact, analysis of other BaseScope samples point to different overall dynamics, especially with respect to sub-clone replicative advantage, *s* (Figs S24-S31).

The observed distributions of inferred values for mutant replicative advantage, *s*, and cell pushing, *q*, within the same BaseScope image could be due to a number of reasons. The experimental tissue represents 2-dimensional samples taken from a 3-dimensional system along a randomly oriented plane. As previously discussed, this sampling can have significant impacts on the appearance of the sub-clonal pattern, potentially creating the false appearance of spatially distinct sub-clones. When analysed individually, these small isolated sub-clones are likely to lead to variances in the inferred dynamics. Beyond this, however, such variances in inferred parameter values could reflect local fluctuations in the dynamics of the system. Cell mixing is likely to be a result of complex combination of cell-intrinsic and extrinsic factors, influenced by physical stresses exerted by the tumour microenvironment. Physical stresses may fluctuate even across small areas of tissue, and inferring fluctuating values of cell pushing in different areas of the tumour may be consistent with this notion. In line with this reasoning, the observed distribution of mutant replicative advantage, *s*, could reflect the varying propensity of mutant cells to proliferate, which can be expected to fluctuate spatially. Whilst certain genetic alterations may confer an cell-intrinsic increase in proliferation rate, other cell-extrinsic factors are also expected to contribute to the “effective” replication rate of any particular cell, such as the availability of free space, access to nutrients, and proximity to the immune system.

## Discussion

The difficulty of obtaining longitudinal samples in cancer studies calls for novel methods to analyse single time sampled data, especially spatial information, to unravel early tumour dynamics *in vivo*. Spatially resolved tumour samples have been studied in a multitude of ways in the prognosis of cancer, yet spatial information may also have the potential to inform on the past sub-clonal dynamics. Here, we leveraged statistics of random walkers as a new methodology to quantify complex patterns of sub-clonal mixing observed *in vivo*. We first characterised the normalised class mean first passage time (CMFPT) using a set of standard two-class toy model patterns. By normalising each image measurement to that from a null model, in which the image pixels are rearranged uniformly at random, we were able to compare CMFPT data for patterns of different total sizes, and different ratios of the two classes. These simple toy model measurements demonstrated the power of the CMFPT to distinguish between patterns of different class ratios and underlying pattern structures. Patterns with weak and strong segregation of the two classes were represented in radically different regions of the phase space.

Extending our analysis to a large dataset of simulated sub-clonal mixing patterns generated with our agent-based model, we showed that the CMFPT is capable of distinguishing between different regimes of sub-clonal dynamics, through the resulting patterns of sub-clonal mixing. Tumours with weakly advantageous mutant sub-clones, which emerged late during the wild-type (WT) population expansion, led to spatial patterns with vastly different first passage times to those with more rapidly replicating mutant sub-clones, those with relatively early emerging sub-clones, or both. Consistent with other recent computational modelling based on genomic sequencing data [4, 33], we find that earlier arising mutant sub-clones lead to greater variegation in the final tumour. Whilst these effects have been studied through multiomic sequencing analysis of spatial bulk sub-samples, here we show for the first time that these dynamics can also be measured using the CMFPT applied to cellular resolution maps of the sub-clonal architecture.

These analyses demonstrate how the CMFPT provides a more detailed description of the patterns of mutant sub-clonal mixing than the spatial Shannon’s entropy measure that was previously used to analyse the BaseScope patterns. Whilst the agent-based model used in this study and the original BaseScope study were similar, here we also explored the influence of varying the strength of cell pushing. We showed that cell pushing can profoundly affect the topology of the mutant sub-clonal pattern. Whilst the patterns produced under a surface growth model and one with cell pushing were qualitatively similar when analysed using Shannon’s entropy, such dynamics were measurable when analysing the mixing patterns at multiple spatial scales using the CMFPT.

By applying our CMFPT method to human colorectal cancer samples analysed with BaseScope, we compared statistical features of the experimental patterns to the simulated sub-clonal patterns. In doing so we inferred values for the replicative advantage of sub-clonal cells, *s*, the relative time at which the sub-clone infiltrated the local WT population, *t*_*mut*_, and the strength of cell pushing, *q*. We uncovered a wide range of dynamics, with some samples consistent with weak mutant selection, and others suggesting more aggressively expanding sub-clones. We predicted that mutant sub-clones detected with BaseScope mostly arose early in the expansion of the WT population, and that cell pushing was generally weak *in vivo*. By analysing multiple sub-regions of the sub-clonal patterns in BaseScope samples 02, 28 and 34 we were able to obtain a more detailed insight into the sub-clonal dynamics in these tumours. When viewed in aggregate, the measurements within these samples formed a trend suggestive of highly aggressive mutant sub-clones, with rates of replication as high as 3 or 4 times that of the WT population. Whilst we were only able to infer local values for the relative time of sub-clonal invasion, the trend formed by these measurements allows us to speculate at the time of the original mutational event in the tumours. For instance, BaseScope sample 28 contains a greater fraction of sub-regions which are best described by late invading sub-clones compared to samples 02 and 34. This suggests that the original mutational event occurred later in the evolution of this tumour than those in samples 02 and 34. To make a more qualitative statement would require extensive 3-dimensional modelling and remains a challenge for future work. However, our analysis of these samples demonstrates how important the role of sampling is in quantifying tumour evolution and how additional dynamics can be revealed by comparing sub-regions within the same tumour.

Alongside the measured sub-clonal patterns published in the original paper on BaseScope (Ref [20]) we also applied our method to a small number of colorectal cancer samples published in a recent study by Heide *et al*. [33]. In their work, Heide and colleagues combine multi-region sequencing data of colorectal tumours with spatial computational modelling to infer values for sub-clonal selection, relative time of sub-clone invasion and cell pushing strength. Samples from two patients in this study were also analysed with BaseScope (samples A7, A10 & A11 from patient C537, and A12 from patient C539; Fig S1a), giving us an opportunity to compare the modelling framework of Heide and colleagues to our orthogonal approach of analysing patterns of sub-clonal mixing to study the evolution of these tumours. For instance, positive sub-clonal selection was predicted by the model of Heide *et al*. in patient C539. Analysing BaseScope sample A12 from the same patient using CMFPT, we also inferred some moderate positive sub-clonal selection (indicated by *s* = 0.2), and a moderately early relative time of sub-clone invasion (*t*_*mut*_ = 0.08) which is broadly consistent with the predictions made by Heide and colleagues.

Our predictions for patient C537, however, deviate from those made by Heide *et al*.. Whilst they predicted neutral sub-clonal evolution in this tumour, our modelling suggested strong sub-clonal selection (best-fit values of *s* = 3, 0.2 & 3 for samples A7, A10 & A11 respectively). The variations of predicted sub-clonal selection strength within samples A7, A10 & A11 using our method, and the disparity between our predictions and those made by Heide *et al*., highlight the challenge of dealing with the stochasticity of the tissue sampling process when analysing BaseScope patterns. Whilst 3-dimensional tumour models have been studied in the past [6, 34], and our simulations can be extended from 2 to 3-dimensions, our primary focus was on developing the methodology of CMFPT measurements. Future research into the statistics of random spatial sampling of 3-dimensional sub-clonal patterns could help mitigate uncertainty associated with “random” spatial sampling of real tumour tissue.

Our simulation model accounts for two clinically relevant factors: the strength of sub-clonal selection (*s*) and the relative time of mutant sub-clone infiltration in the WT population (*t*_*mut*_). While translating less straight-forwardly to *in vivo* systems, our third model parameter, cell pushing strength (*q*), was shown to dramatically affect the resulting sub-clonal mixing patterns. Using a different modelling approach, Ryser *et al*. recently showed that short range cell migration may also have important implications for sub-clonal mixing in colorectal cancers [8], especially in the early stages of tumourigenesis, and cell migration has been studied in other computational models of tumour growth [6, 35, 36]. Where Ryser *et al*. found early tumour cell mobility to increase sub-clonal mixing in the final tumour, we find that strong cell pushing strength produces similar results.

In summary, we have shown that the CMFPT can be used to quantify clustering and heterogeneity of sub-clonal patterns in spatially resolved tumour samples. We extended this method from previous applications in spatial demographic data to patterns of sub-clonal mixing in colorectal tumour samples acquired with BaseScope, and combined experimental measurements with spatial computational modelling of expanding tumour populations to estimate parameters of early sub-clonal evolution in our samples. In line with contemporary models of tumour evolution, our analysis suggests that the large tumour sub-clones which were observed may have arisen early in the expansion of the wild-type tumour cell population, exhibiting a range of selective growth advantages over wild-type cells. Our work lays a foundation for understanding early cancer evolution from high resolution spatial data. Further research to determine the influence of 2-dimensional sampling on inference of the dynamics, and effectively modelling the role of the tumour microenvironment, is required to further elucidate early cancer dynamics.

## Acknowledgements

M.J.H. is supported by the Life Sciences Institute at Queen Mary University of London. A.B. acknowledges funding from the Juan de la Cierva program; the Spanish Ministry of Universities; the European Union - Next Generation EU; the Recovery, Transformation and Resilience Plan; the University of the Balearic Islands; the Departament d’Enginyeria Informàtica i Matemàtiques, Universitat Rovira i Virgili, Tarragona, Spain, and Instituto de Física Interdisciplinar y Sistemas Complejos IFISC (CSIC-UIB), Campus UIB, 07122 Palma de Mallorca, Spain.

## Methods

### Code availability

Scripts designed to generate and visualise sub-clonal mixing patterns are available at: https://github.com/MagnusHaughey/CancerSubclonalPatterns. Code relating to calculations of first passage time statistics can be found at: https://mygit.katolaz.net/covid_19_ethnicity/rw-segregation.

### Simulating random walkers and estimating first passage time statistics

To assess the spatial heterogeneity of cell types we use the normalised mean first passage times 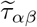 between class *α* and class *β* [25]. The quantity 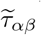 is obtained by dividing the mean first passage time, the average number of steps *τ*_*αβ*_ taken by a walker to arrive at a node of class *β* when departing from a node of class *α*, by its null-model counterpart, 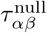. To estimate 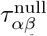 we simulate random walks on a graph with the same connectivity as the original graph, but with the node classes redistributed uniformly at random, such that for an original graph with class ratio *ϕ* = *x*, any arbitrarily selected coloured node will be of class *α* and *β* with probabilities *P*_*α*_ = *x* and *P*_*β*_ = 1 − *x* respectively. Since the quantity 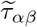 is normalised by a null-model, it is dimensionless and represents whether the time to arrive at a node of a given class is smaller or larger than the corresponding transition in the null-model. For example, if 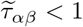 then the expected number of steps taken before arriving at a node of class *β* is less in the experimental pattern than in the null-model, and is greater when 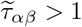.

To reduce computational time, we downsample the BaseScope data by binning image pixels. When altering the image resolution, the proportion of pixel classes is preserved in each bin. For example, if the pixels falling into a particular bin consisted of 50% red and 50% yellow pixels, then a walker on the downsampled network will encounter either a red or yellow bin at these coordinates with equal probabilities of 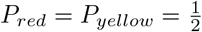. We downsample all BaseScope and simulated sub-clonal patterns to the same resolution, 3000 pixels in total, prior to analysis.

### Spatial simulations of sub-clonal evolution in tumours

We generate cell mixing patterns using the Gillespie algorithm [37] to simulating stochastic birth and death of cells on a 2-dimensional square lattice. Since the BaseScope assay targets and detects specific genetic point mutations, we adopt a binary system in which a particular cell can either be wild-type (WT) or mutated. A simulation begins by seeding a single WT cell on the lattice, which has a birth rate greater than its death rate, and terminates when the entire system (WT and mutated cells) reaches a specified size *N*_*max*_. During initial growth, the WT clone expands until it reaches a size *n* = *N*_*max*_ · *t*_*mut*_ cells, at which point an existing cell is randomly chosen to acquire a mutation. All subsequent cells related to the newly mutated cell will inherit the mutation, and cells can not revert from the mutated state back to WT.

Cells carrying the mutation experience a relative fitness advantage compared to WT cells. This fitness advantage manifests as an increased birth rate, giving the birth rate of cell *i, b*_*i*_, as

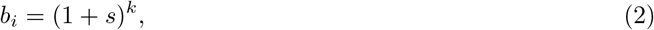

where *k* ∈ {0, 1}, and *s* represents the relative fitness advantage conferred by the mutation. Cell death is coupled to cell birth in the simulations. Before a cell divides, it is randomly determined either to continue with the division, or die giving a death rate for cell *i, d*_*i*_, of

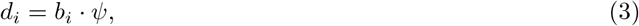

where *ψ* ∈ [0, 1], and is fixed at *ψ* = 0.3 throughout this study unless stated otherwise. When a cell dies it immediately is removed from the lattice leaving behind an empty lattice point.

To model the displacement of neighbouring cells during division, we adapted an algorithm developed by Waclaw et al. [6], which involves the dividing cell searching its neighbourhood and finding a “path” to one of the nearest empty lattice points. In order to create space to divide into two daughter cells, the dividing cell pushes neighbouring cells along this path, filling the nearby empty lattice point and creating a new empty space adjacent to itself. In our simulations we implement these mechanics and extend them by accounting for relative positions of cells during pushing, favouring straight-line pushing over displacement in other directions, in accordance with Newton’s second law of motion. The parameter *q* determines the maximum allowed size for the constructed path in this algorithm. If no path can be found which is less than or equal to *q* in length, the cell cannot successfully divide. Setting this parameter to *q* = 0 leads to surface growth dynamics, in which a cell must be adjacent to an empty lattice point in order to divide successfully.

Model parameters used to generate *in silico* mixing patterns are listed in Table S1. Note that, due to decreased probability of sub-clonal survival for parameter combinations (*s, t*_*mut*_, *q*) = (0, 0.5, 20), (0.1, 0.5, 20) and (0.2, 0.5, 20), sub-clonal pattern data could not be generated for these parameter combinations.

### Bayesian grid-search for best-fit model parameter inference

To perform inference of mutant replicative advantage (*s*), mutation timing (*t*_*mut*_) and cell pushing strength (*q*) for each BaseScope sub-sampled region (Figs S8-S23), we implement a Bayesian-style grid search. For each experimental pattern, we sample the posterior distribution of inferred model parameters *s, t*_*mut*_ and *q* by finding the *n* nearest simulated sub-clonal patterns in the 4-dimensional phase space of class mean first passage times 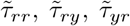 and 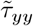, as determined by log-Euclidean distance. We chose a posterior sample size of *n* = 250 for each experimental pattern, and median as a point estimate. To select the “best fit” simulated sub-clonal pattern we found the pattern generated with the median parameter values within the posterior samples that minimised the log-Euclidean distance to the experimental image (marginal median method). For a small number of experimental samples (BaseScope sample 11, sub-sample 1; sample 17a2, sub-sample 1; sample 28, sub-samples 5&11; sample 34, sub-sample 2), there was no posterior sample that corresponded to the inferred median parameter values. This was due to these experimental samples falling between multiple clusters of simulated patterns that corresponded to very early and very late arising mutant populations, resulting in the posterior distribution consisting of samples from both clusters. In these cases, the median was not a suitable point estimator of the “best fit” model parameter values. Instead, the most abundant combination of *s, t*_*mut*_ and *q* within the posterior samples was chosen as the “best-fit” parameters (3D histogram method). Table S2 lists the best-fit values of *s, t*_*mut*_ and *q* as determined using the marginal median and 3D histogram methods.

## Notes

### Competing Interest Statement

The authors have declared no competing interest.

