## Supplemental Information for "First passage time analysis of spatial mutation patterns reveals evolutionary dynamics of pre-existing resistance in colorectal cancer"

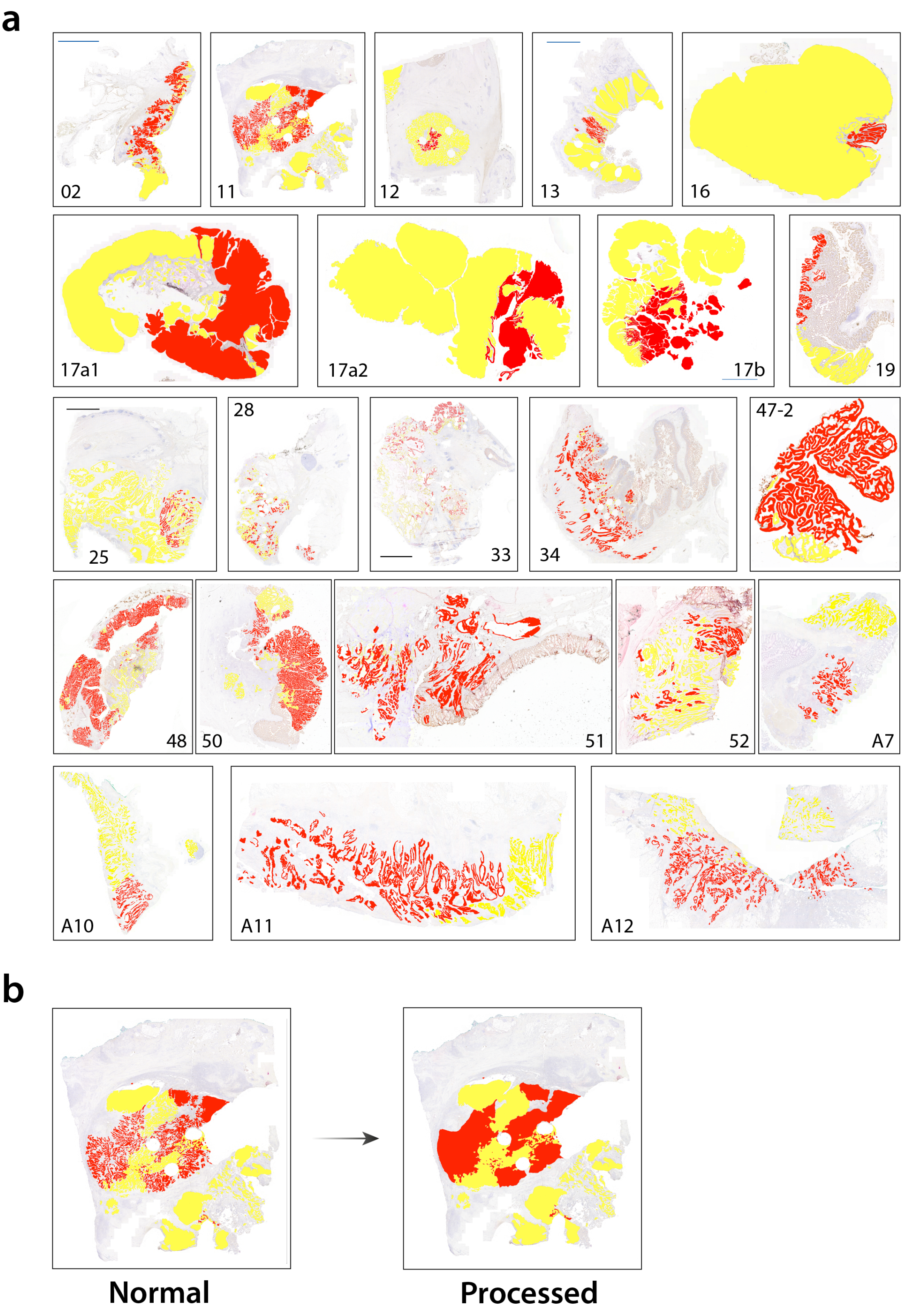

**Fig S1. All BaseScope patterns analysed in this study.** **(a)** Sub-clonal mutations in colorectal tumours analysed with BaseScope used in this study, from Refs [1, 2]. Colorectal tumour cells and surrounding epithelium is pictured, with wild-type tumour cells highlighted in yellow, and mutated sub-clonal tumour population highlighted in red. Non-cancerous tissue is not highlighted. **(b)** Representative example of a "raw" BaseScope image, and the same image after application of the pre-processing steps described in the main text.

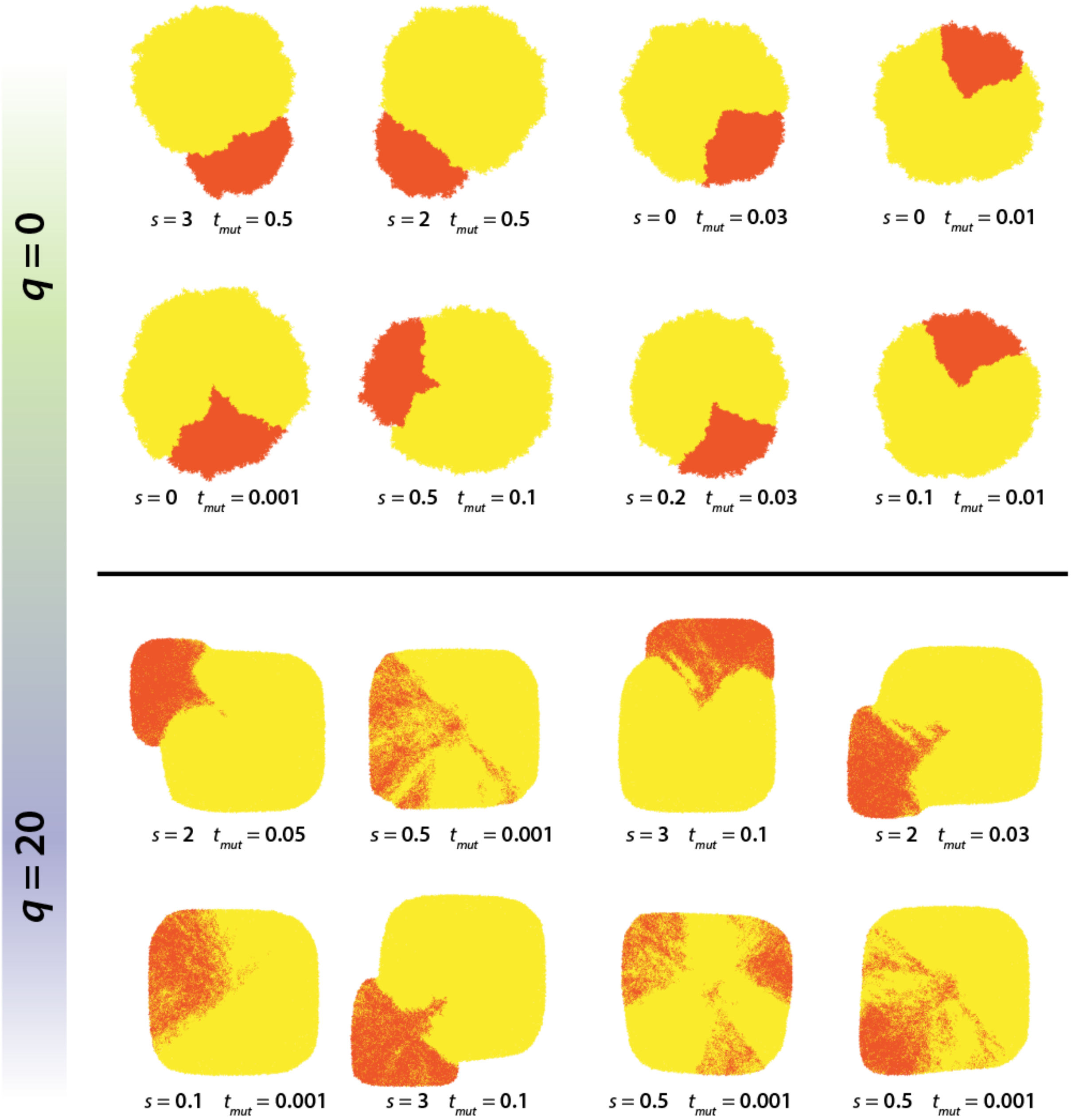

**Fig S2. Diversity of simulated sub-clonal patterns.** Representative sub-clonal mixing patterns, each with class ratio $\approx 0.2$, produced with the spatial computational model for pushing strengths $q = 0$ and $q = 20$. A lower pushing values the diversity in patterns is smaller, for constant class ratio. As pushing strength is increased, patterns with the same class ratio are more diverse.

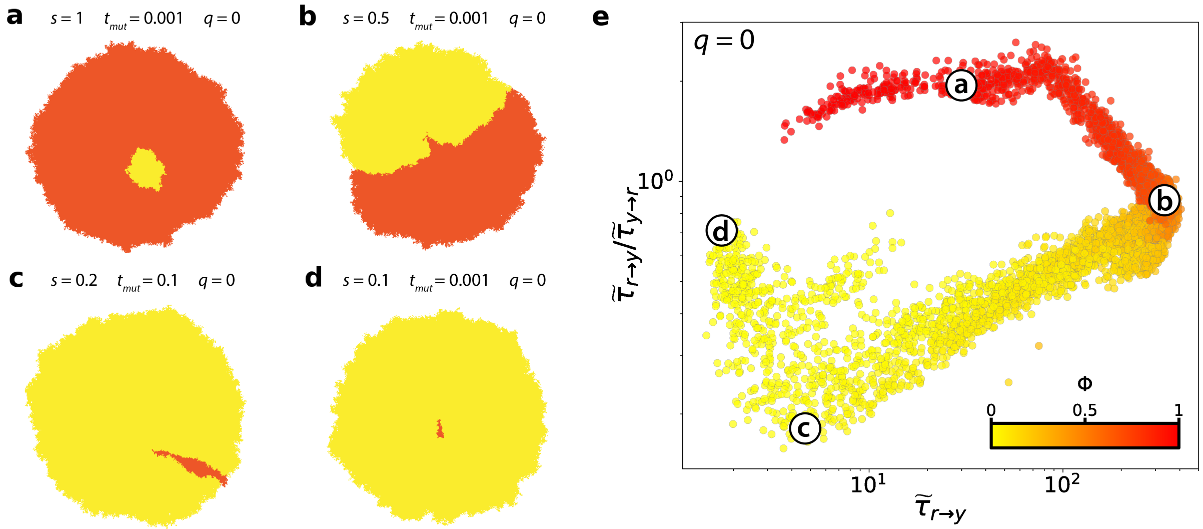

**Fig S3. Analysis of simulated sub-clonal patterns. (a-d)** Representative examples of tumour sub-clonal patterns simulated with a pushing strength of $q = 0$. **(e)** CMFPT analysis of all simulated sub-clonal patterns with a pushing strength of $q = 0$ in the $(\widetilde{\tau}_{ry},\widetilde{\tau}_{ry}/\widetilde{\tau}_{yr})$ phase space. Points are coloured according to pattern class ratio, $\phi$, and images shown in (a-d) are highlighted in the phase space.

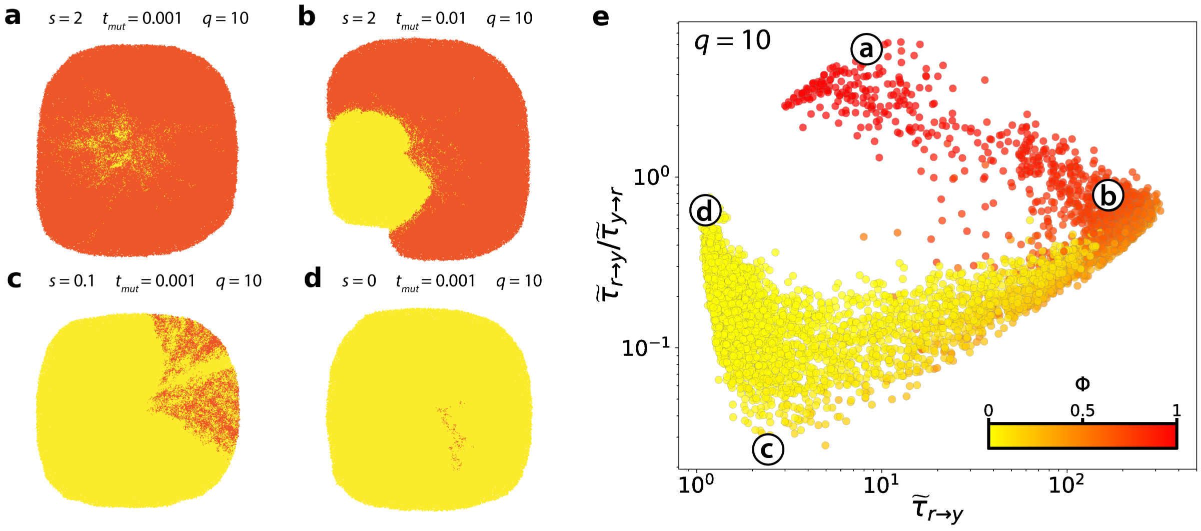

**Fig S4. Analysis of simulated sub-clonal patterns. (a-d)** Representative examples of tumour sub-clonal patterns simulated with a pushing strength of $q = 10$. **(e)** CMFPT analysis of all simulated sub-clonal patterns with a pushing strength of $q = 10$ in the $(\widetilde{\tau}_{ry}, \widetilde{\tau}_{ry}/\widetilde{\tau}_{yr})$ phase space. Points are coloured according to pattern class ratio, $\phi$, and images shown in (a-d) are highlighted in the phase space.

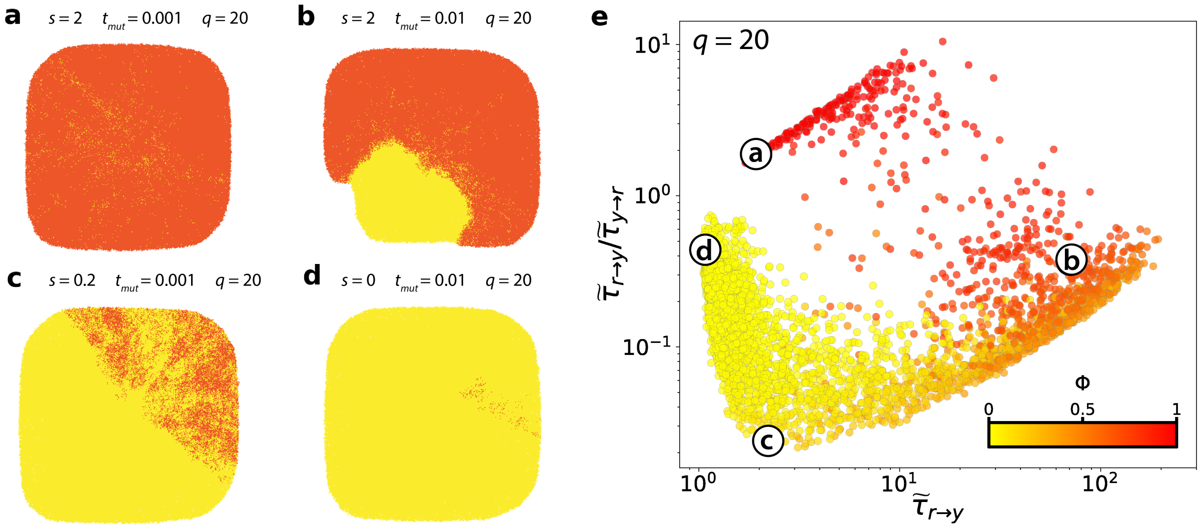

**Fig S5. Analysis of simulated sub-clonal patterns. (a-d)** Representative examples of tumour sub-clonal patterns simulated with a pushing strength of $q = 20$. **(e)** CMFPT analysis of all simulated sub-clonal patterns with a pushing strength of $q = 20$ in the ($\widetilde{\tau}_{ry}$,$\widetilde{\tau}_{ry}/\widetilde{\tau}_{yr}$) phase space. Points are coloured according to pattern class ratio, $\phi$, and images shown in (a-d) are highlighted in the phase space.

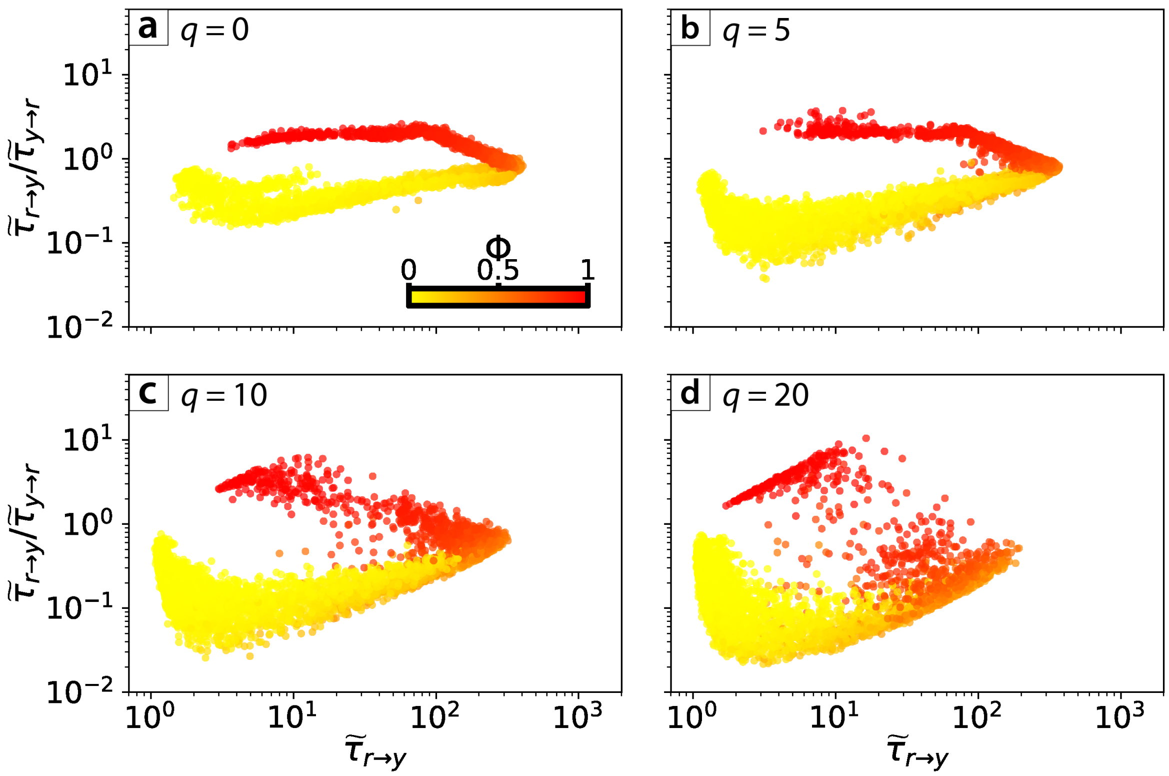

**Fig S6. Analysis of simulated sub-clonal patterns for varying model parameter values.** Images generated by the model in the phase space ($\tilde{\tau}_{ry}$,$\tilde{\tau}_{ry}/\tilde{\tau}_{yr}$), with points coloured according to values of model parameter $s$ and point size depending on parameter $t_{mut}$. Images are separated depending on their pushing value **(a)** 0, **(b)** 5, **(c)** 10 and **(d)** 20.

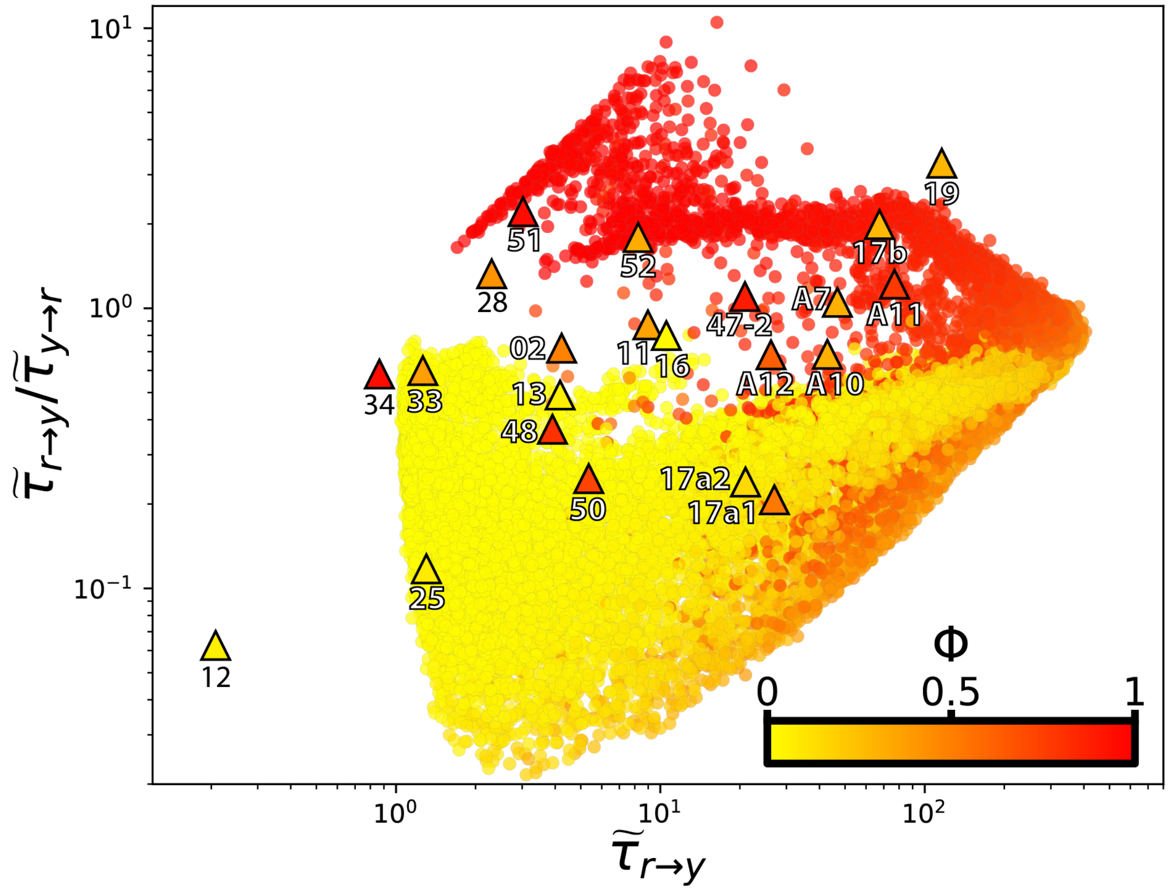

**Fig S7. Initial CMFPT analysis of sub-clonal patterns in human colorectal tumour samples.** CMFPT analysis of human colorectal tumour samples plotted in the ($\widetilde{\tau}_{ry}$,$\widetilde{\tau}_{ry}/\widetilde{\tau}_{yr}$) phase space. Each triangular marker represents a single colorectal cancer sample analysed with BaseScope. Circular points represent all simulated sub-clonal patterns, and points are coloured according to pattern class ratio, $\phi$.

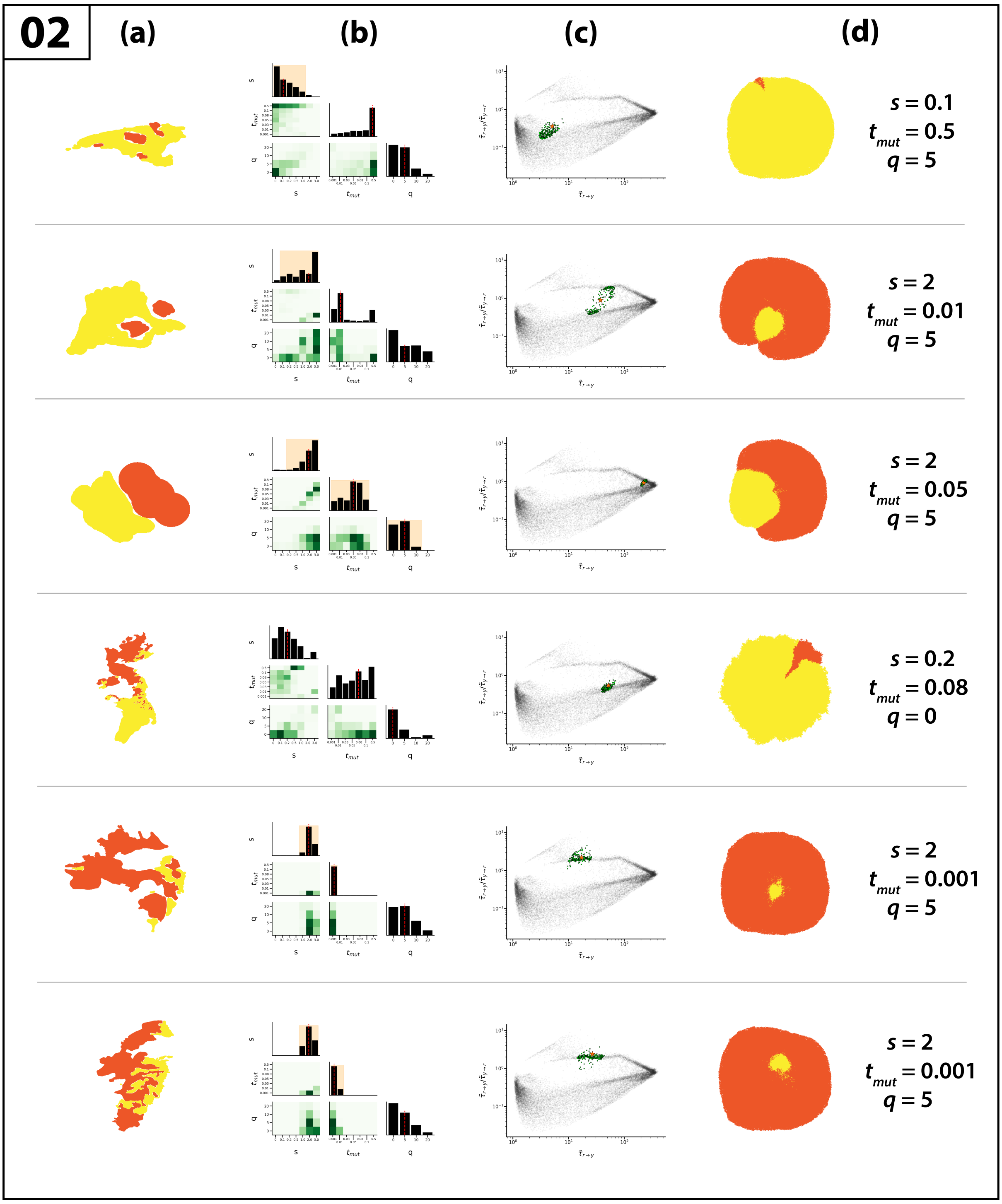

**Fig S8.** Bayesian grid-search analysis of BaseScope sample 02. **(a)** Sub-sample of sample 02. **(b)** Marginal posterior distributions of model parameters $s$, $t_{mut}$, and $q$, representing mutant selection strength, mutation timing and cell pushing strength respectively. Median inferred parameter value is indicated by the vertical dashed line along the diagonal panels. 95% credible regions lie within the shaded region in the diagonal panels. Where no shaded region is given, this interval was the entire parameter range. **(c)** All analysed simulated sub-clonal mixing patterns (grey points) with CMFPT value of the BaseScope sub-sample (star) and posterior samples (green points). **(d)** Best-fit simulated sub-clonal pattern and parameters representing the most abundant parameter combination within the posterior distribution.

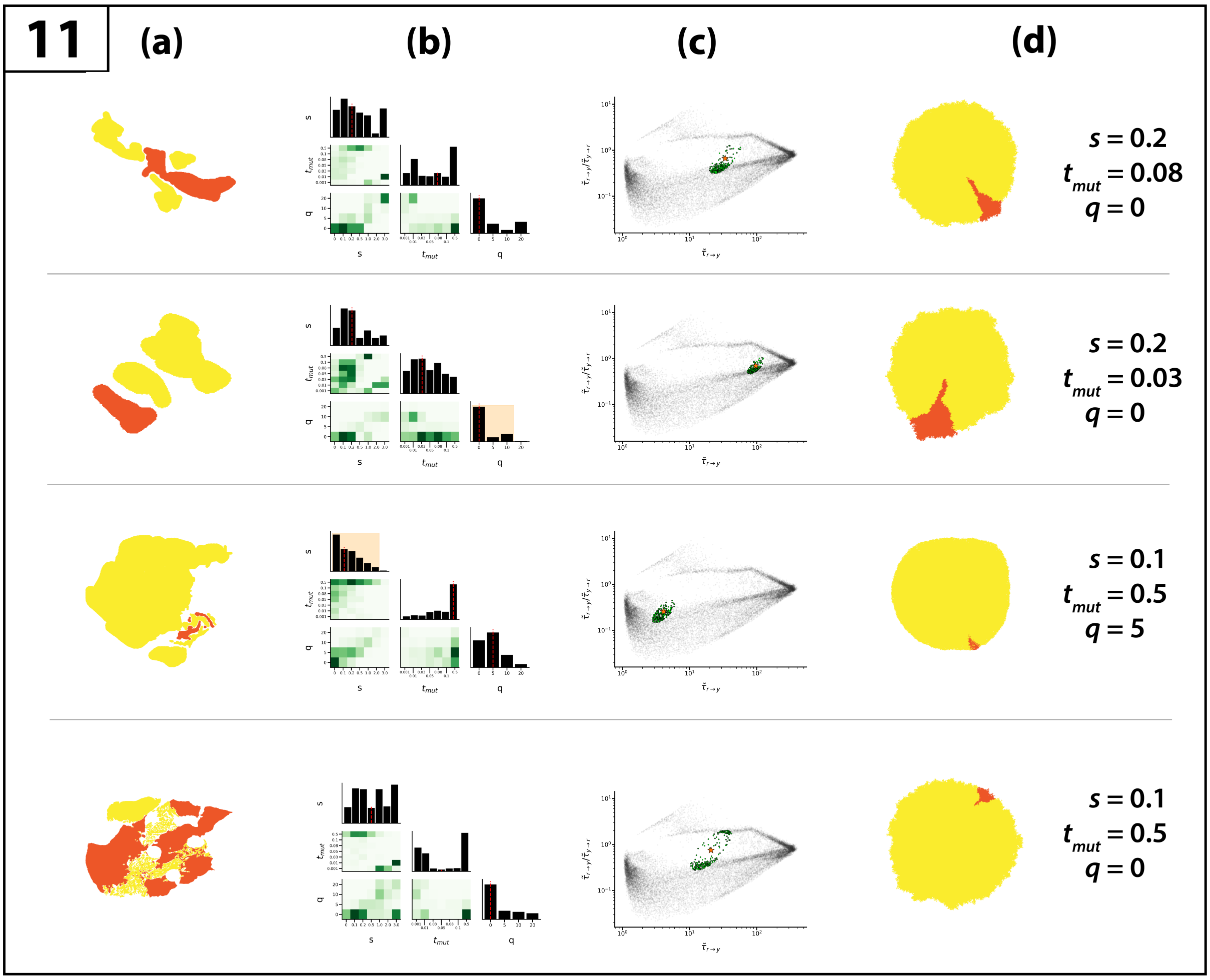

**Fig S9.** Bayesian grid-search analysis of BaseScope sample 11. **(a)** Sub-sample of sample 11. **(b)** Marginal posterior distributions of model parameters $s$, $t_{mut}$, and $q$, representing mutant selection strength, mutation timing and cell pushing strength respectively. Median inferred parameter value is indicated by the vertical dashed line along the diagonal panels. 95% credible regions lie within the shaded region in the diagonal panels. Where no shaded region is given, this interval was the entire parameter range. **(c)** All analysed simulated sub-clonal mixing patterns (grey points) with CMFPT value of the BaseScope sub-sample (star) and posterior samples (green points). **(d)** Best-fit simulated sub-clonal pattern and parameters representing the most abundant parameter combination within the posterior distribution.

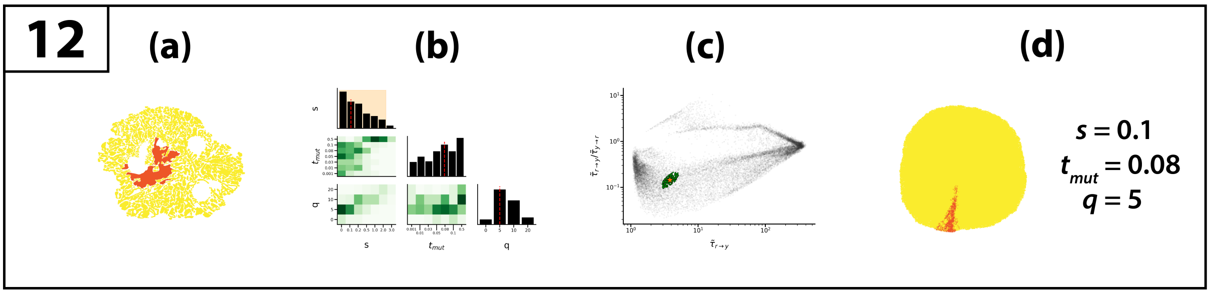

**Fig S10.** Bayesian grid-search analysis of BaseScope sample 12. **(a)** Sub-sample of sample 12. **(b)** Marginal posterior distributions of model parameters $s$, $t_{mut}$, and $q$, representing mutant selection strength, mutation timing and cell pushing strength respectively. Median inferred parameter value is indicated by the vertical dashed line along the diagonal panels. 95% credible regions lie within the shaded region in the diagonal panels. Where no shaded region is given, this interval was the entire parameter range. **(c)** All analysed simulated sub-clonal mixing patterns (grey points) with CMFPT value of the BaseScope sub-sample (star) and posterior samples (green points). **(d)** Best-fit simulated sub-clonal pattern and parameters representing the most abundant parameter combination within the posterior distribution.

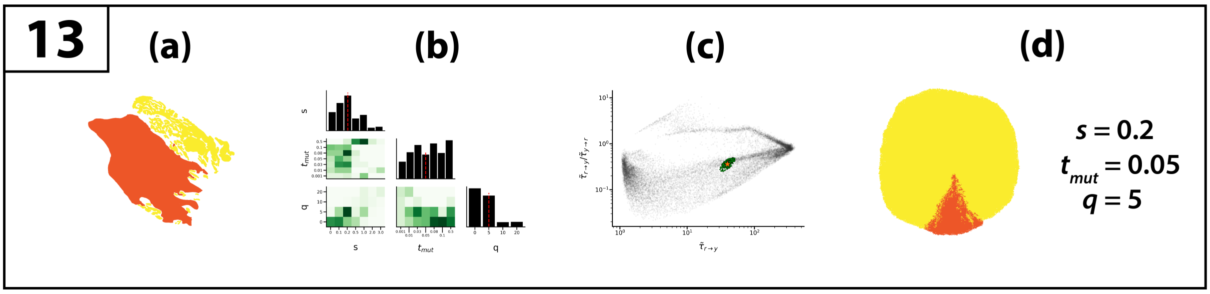

**Fig S11.** Bayesian grid-search analysis of BaseScope sample 13. **(a)** Sub-sample of sample 13. **(b)** Marginal posterior distributions of model parameters $s$, $t_{mut}$, and $q$, representing mutant selection strength, mutation timing and cell pushing strength respectively. Median inferred parameter value is indicated by the vertical dashed line along the diagonal panels. 95% credible regions lie within the shaded region in the diagonal panels. Where no shaded region is given, this interval was the entire parameter range. **(c)** All analysed simulated sub-clonal mixing patterns (grey points) with CMFPT value of the BaseScope sub-sample (star) and posterior samples (green points). **(d)** Best-fit simulated sub-clonal pattern and parameters representing the most abundant parameter combination within the posterior distribution.

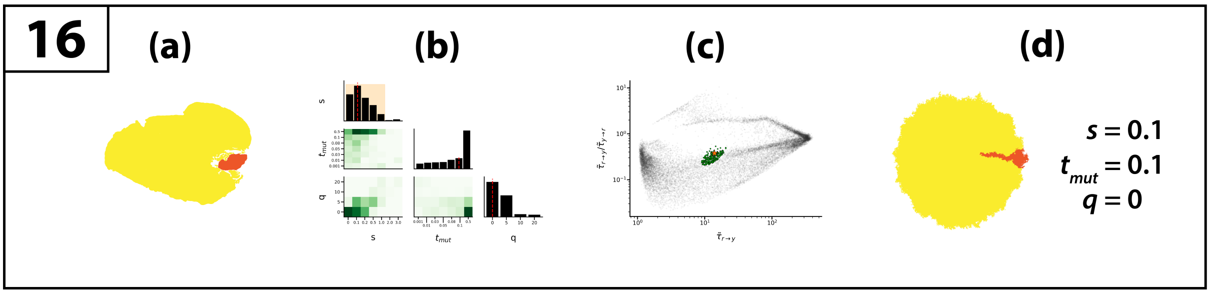

**Fig S12.** Bayesian grid-search analysis of BaseScope sample 16. **(a)** Sub-sample of sample 16. **(b)** Marginal posterior distributions of model parameters $s$, $t_{mut}$, and $q$, representing mutant selection strength, mutation timing and cell pushing strength respectively. Median inferred parameter value is indicated by the vertical dashed line along the diagonal panels. 95% credible regions lie within the shaded region in the diagonal panels. Where no shaded region is given, this interval was the entire parameter range. **(c)** All analysed simulated sub-clonal mixing patterns (grey points) with CMFPT value of the BaseScope sub-sample (star) and posterior samples (green points). **(d)** Best-fit simulated sub-clonal pattern and parameters representing the most abundant parameter combination within the posterior distribution.

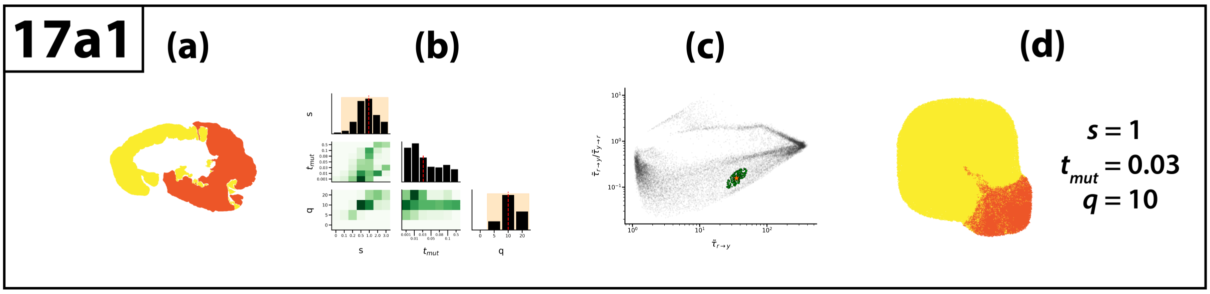

**Fig S13.** Bayesian grid-search analysis of BaseScope sample 17a1. **(a)** Sub-sample of sample 17a1. **(b)** Marginal posterior distributions of model parameters $s$, $t_{mut}$, and $q$, representing mutant selection strength, mutation timing and cell pushing strength respectively. Median inferred parameter value is indicated by the vertical dashed line along the diagonal panels. 95% credible regions lie within the shaded region in the diagonal panels. Where no shaded region is given, this interval was the entire parameter range. **(c)** All analysed simulated sub-clonal mixing patterns (grey points) with CMFPT value of the BaseScope sub-sample (star) and posterior samples (green points). **(d)** Best-fit simulated sub-clonal pattern and parameters representing the most abundant parameter combination within the posterior distribution.

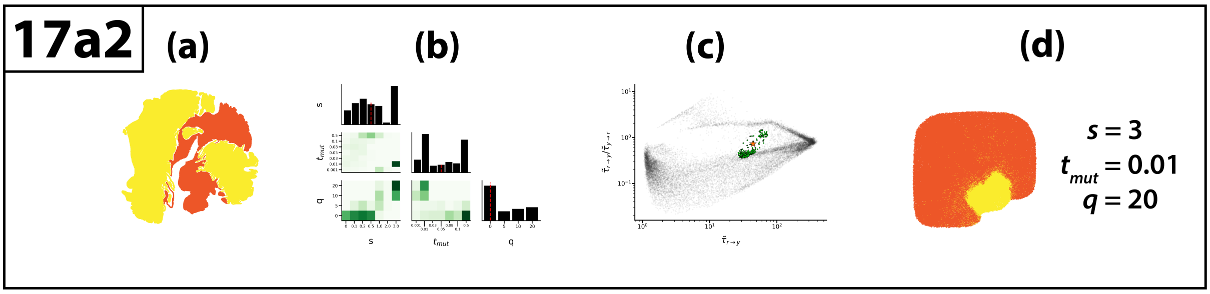

**Fig S14.** Bayesian grid-search analysis of BaseScope sample 17a2. **(a)** Sub-sample of sample 17a2. **(b)** Marginal posterior distributions of model parameters $s$, $t_{mut}$, and $q$, representing mutant selection strength, mutation timing and cell pushing strength respectively. Median inferred parameter value is indicated by the vertical dashed line along the diagonal panels. 95% credible regions lie within the shaded region in the diagonal panels. Where no shaded region is given, this interval was the entire parameter range. **(c)** All analysed simulated sub-clonal mixing patterns (grey points) with CMFPT value of the BaseScope sub-sample (star) and posterior samples (green points). **(d)** Best-fit simulated sub-clonal pattern and parameters representing the most abundant parameter combination within the posterior distribution.

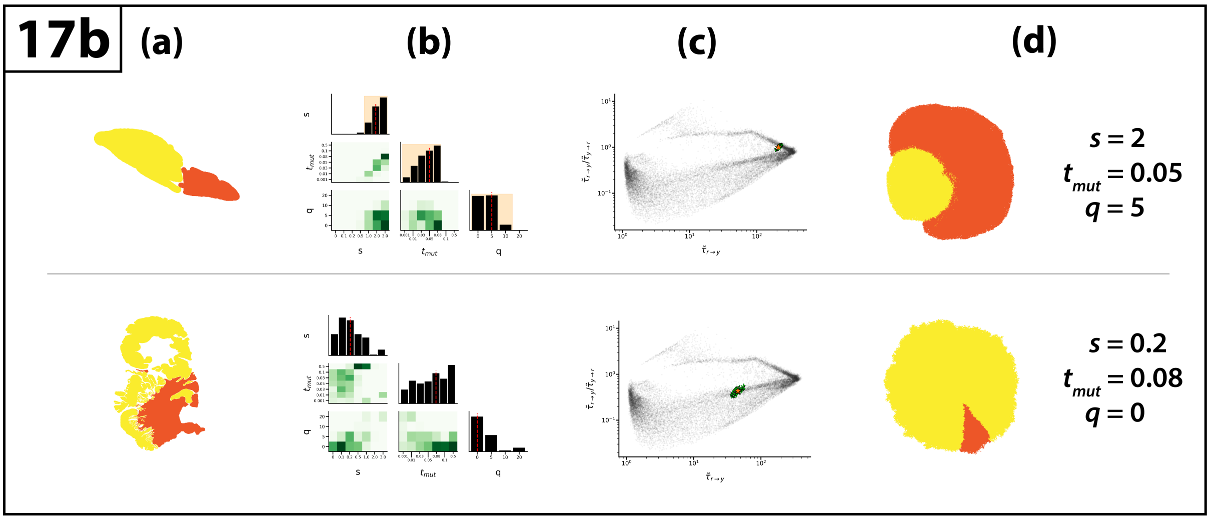

**Fig S15.** Bayesian grid-search analysis of BaseScope sample 17b. **(a)** Sub-sample of sample 17b. **(b)** Marginal posterior distributions of model parameters $s$, $t_{mut}$, and $q$, representing mutant selection strength, mutation timing and cell pushing strength respectively. Median inferred parameter value is indicated by the vertical dashed line along the diagonal panels. 95% credible regions lie within the shaded region in the diagonal panels. Where no shaded region is given, this interval was the entire parameter range. **(c)** All analysed simulated sub-clonal mixing patterns (grey points) with CMFPT value of the BaseScope sub-sample (star) and posterior samples (green points). **(d)** Best-fit simulated sub-clonal pattern and parameters representing the most abundant parameter combination within the posterior distribution.

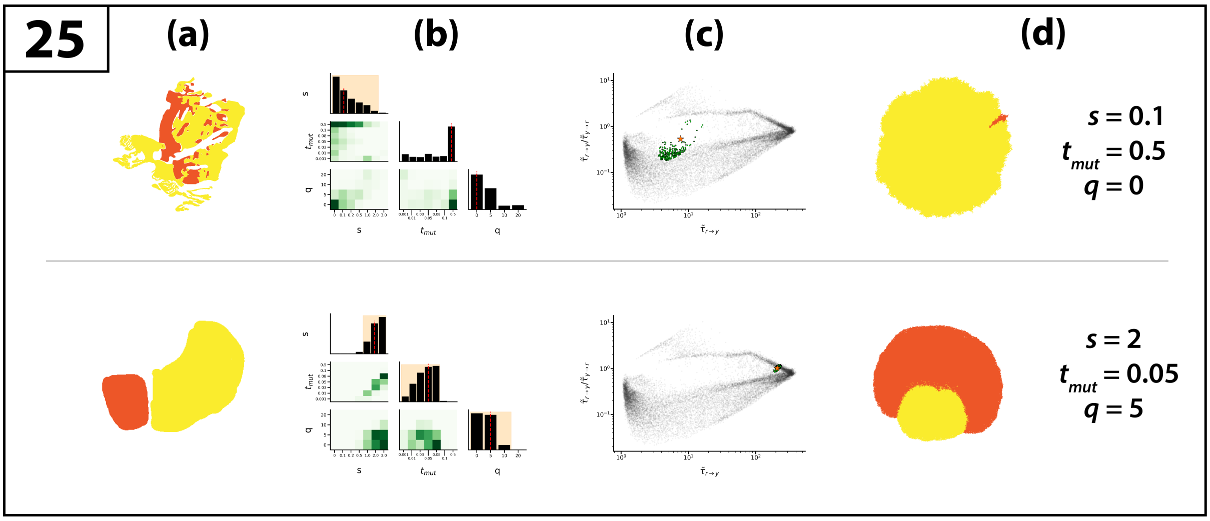

**Fig S16.** Bayesian grid-search analysis of BaseScope sample 25. **(a)** Sub-sample of sample 25. **(b)** Marginal posterior distributions of model parameters $s$, $t_{mut}$, and $q$, representing mutant selection strength, mutation timing and cell pushing strength respectively. Median inferred parameter value is indicated by the vertical dashed line along the diagonal panels. 95% credible regions lie within the shaded region in the diagonal panels. Where no shaded region is given, this interval was the entire parameter range. **(c)** All analysed simulated sub-clonal mixing patterns (grey points) with CMFPT value of the BaseScope sub-sample (star) and posterior samples (green points). **(d)** Best-fit simulated sub-clonal pattern and parameters representing the most abundant parameter combination within the posterior distribution.

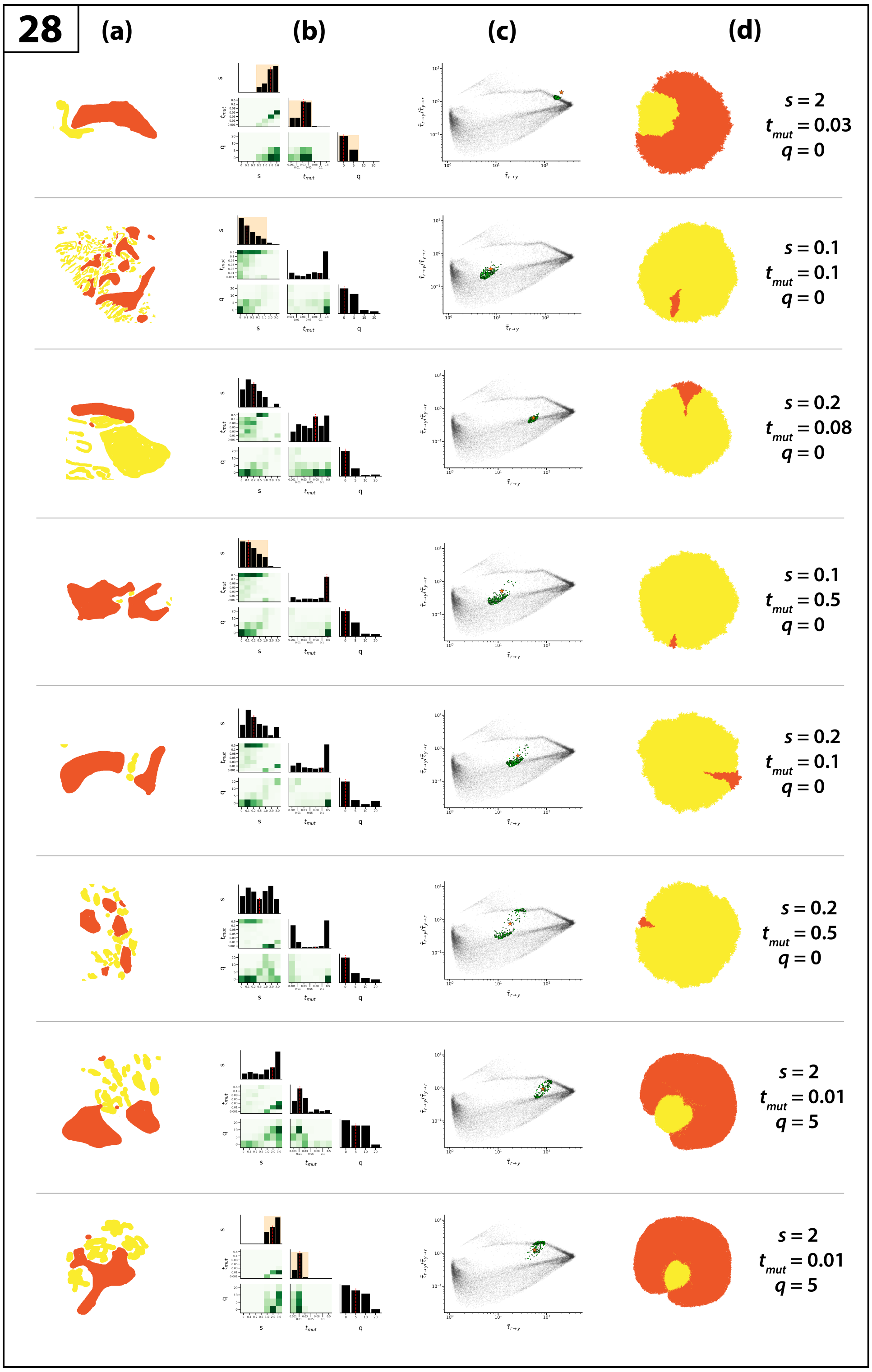

**Fig S17.** Bayesian grid-search analysis of BaseScope sample 28. **(a)** Sub-sample of sample 28. **(b)** Marginal posterior distributions of model parameters $s$, $t_{mut}$, and $q$, representing mutant selection strength, mutation timing and cell pushing strength respectively. Median inferred parameter value is indicated by the vertical dashed line along the diagonal panels. 95% credible regions lie within the shaded region in the diagonal panels. Where no shaded region is given, this interval was the entire parameter range. **(c)** All analysed simulated sub-clonal mixing patterns (grey points) with CMFPT value of the BaseScope sub-sample (star) and posterior samples (green points). **(d)** Best-fit simulated sub-clonal pattern and parameters representing the most abundant parameter combination within the posterior distribution.

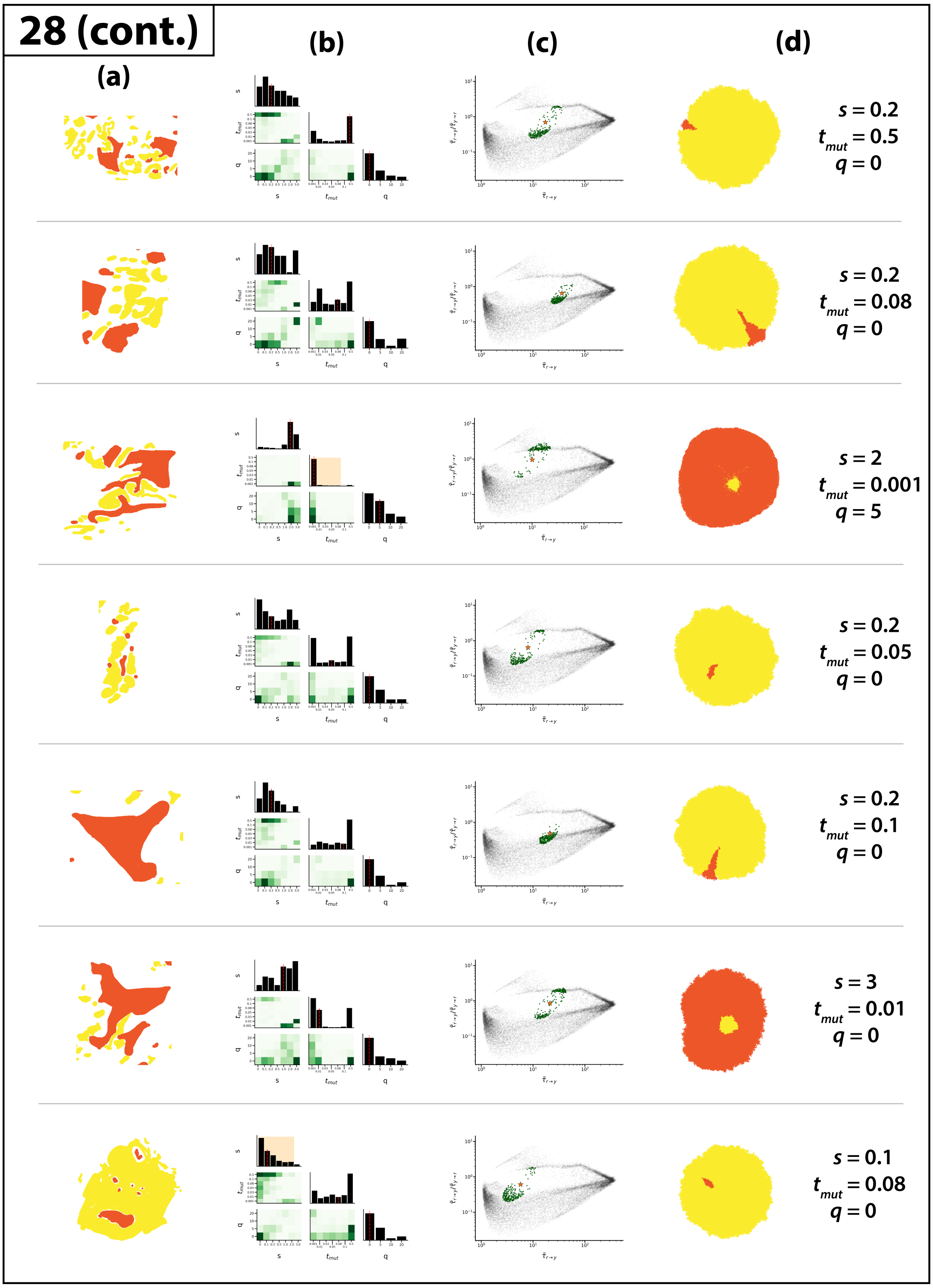

**Fig S18.** Bayesian grid-search analysis of BaseScope sample 28. **(a)** Sub-sample of sample 28. **(b)** Marginal posterior distributions of model parameters $s$, $t_{mut}$, and $q$, representing mutant selection strength, mutation timing and cell pushing strength respectively. Median inferred parameter value is indicated by the vertical dashed line along the diagonal panels. 95% credible regions lie within the shaded region in the diagonal panels. Where no shaded region is given, this interval was the entire parameter range. **(c)** All analysed simulated sub-clonal mixing patterns (grey points) with CMFPT value of the BaseScope sub-sample (star) and posterior samples (green points). **(d)** Best-fit simulated sub-clonal pattern and parameters representing the most abundant parameter combination within the posterior distribution.

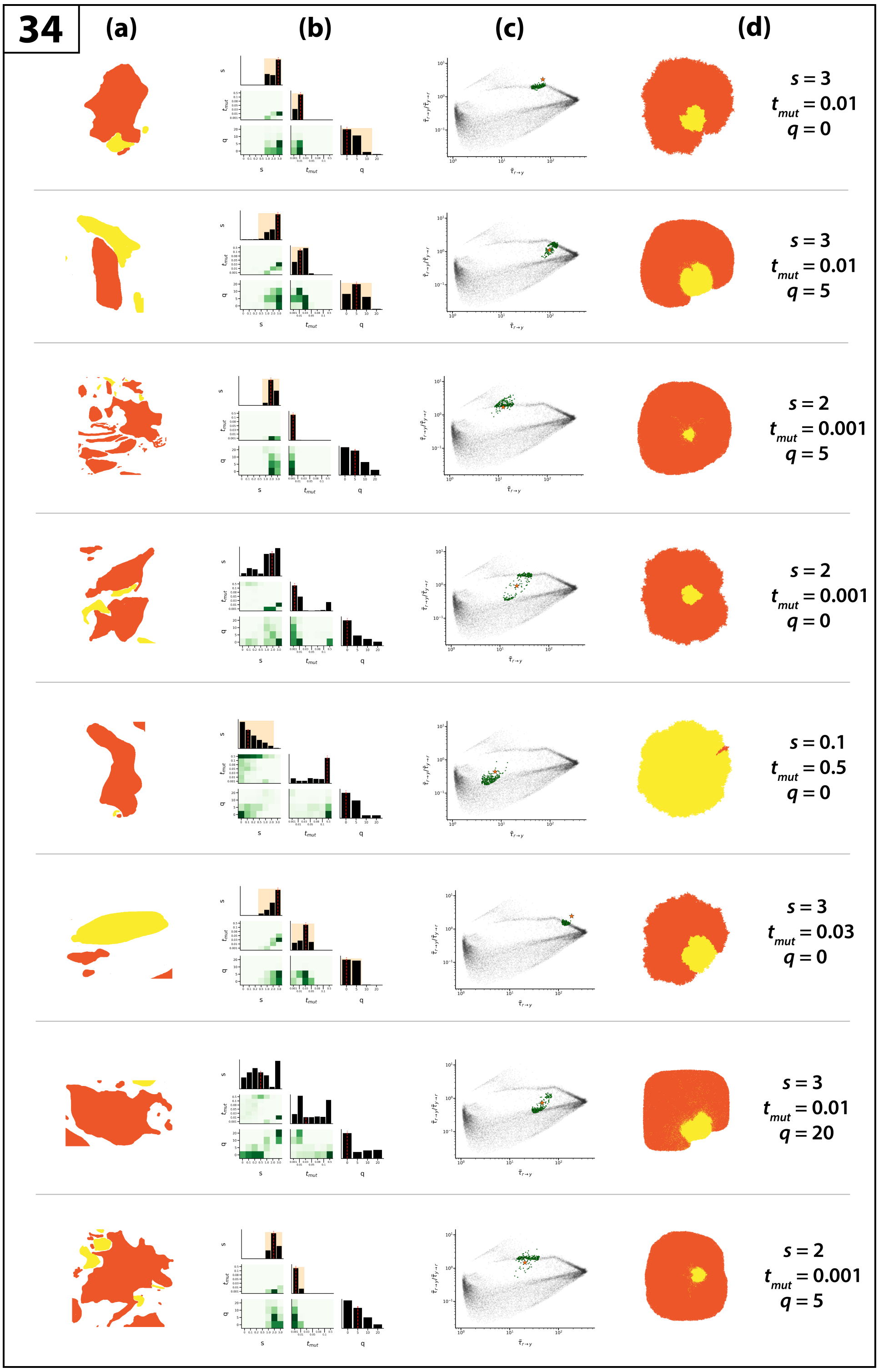

**Fig S19.** Bayesian grid-search analysis of BaseScope sample 34. **(a)** Sub-sample of sample 34. **(b)** Marginal posterior distributions of model parameters $s$, $t_{mut}$, and $q$, representing mutant selection strength, mutation timing and cell pushing strength respectively. Median inferred parameter value is indicated by the vertical dashed line along the diagonal panels. 95% credible regions lie within the shaded region in the diagonal panels. Where no shaded region is given, this interval was the entire parameter range. **(c)** All analysed simulated sub-clonal mixing patterns (grey points) with CMFPT value of the BaseScope sub-sample (star) and posterior samples (green points). **(d)** Best-fit simulated sub-clonal pattern and parameters representing the most abundant parameter combination within the posterior distribution.

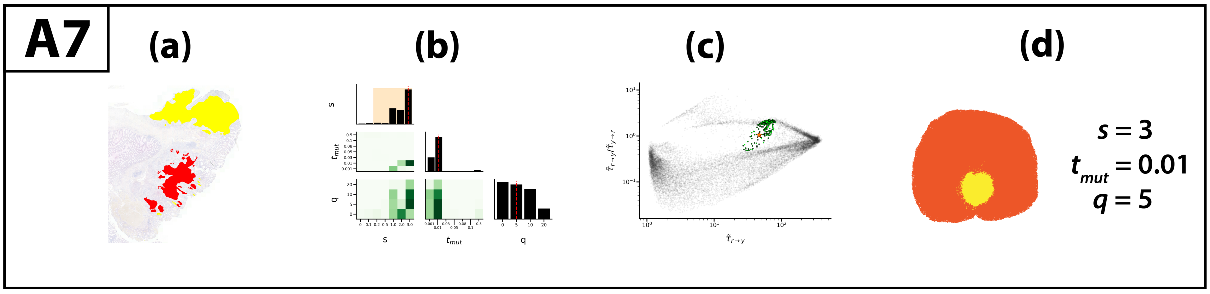

**Fig S20.** Bayesian grid-search analysis of BaseScope sample A7. **(a)** Sub-sample of sample A7. **(b)** Marginal posterior distributions of model parameters $s$, $t_{mut}$, and $q$, representing mutant selection strength, mutation timing and cell pushing strength respectively. Median inferred parameter value is indicated by the vertical dashed line along the diagonal panels. 95% credible regions lie within the shaded region in the diagonal panels. Where no shaded region is given, this interval was the entire parameter range. **(c)** All analysed simulated sub-clonal mixing patterns (grey points) with CMFPT value of the BaseScope sub-sample (star) and posterior samples (green points). **(d)** Best-fit simulated sub-clonal pattern and parameters representing the most abundant parameter combination within the posterior distribution.

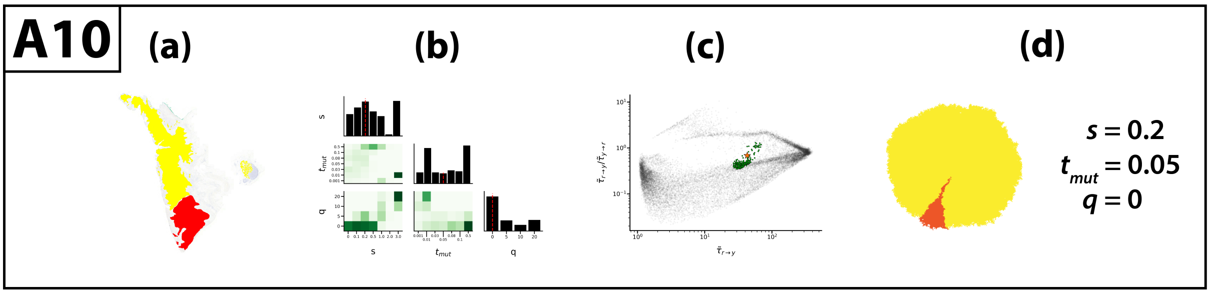

**Fig S21.** Bayesian grid-search analysis of BaseScope sample A10. **(a)** Sub-sample of sample A10. **(b)** Marginal posterior distributions of model parameters $s$, $t_{mut}$, and $q$, representing mutant selection strength, mutation timing and cell pushing strength respectively. Median inferred parameter value is indicated by the vertical dashed line along the diagonal panels. 95% credible regions lie within the shaded region in the diagonal panels. Where no shaded region is given, this interval was the entire parameter range. **(c)** All analysed simulated sub-clonal mixing patterns (grey points) with CMFPT value of the BaseScope sub-sample (star) and posterior samples (green points). **(d)** Best-fit simulated sub-clonal pattern and parameters representing the most abundant parameter combination within the posterior distribution.

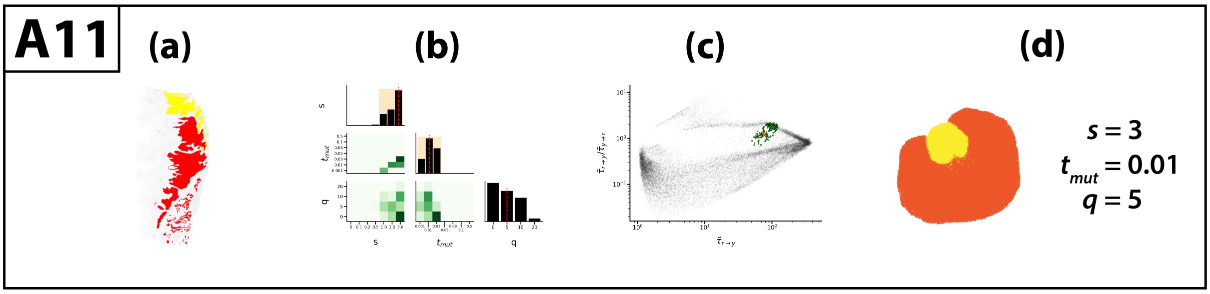

**Fig S22.** Bayesian grid-search analysis of BaseScope sample A11. **(a)** Sub-sample of sample A11. **(b)** Marginal posterior distributions of model parameters $s$, $t_{mut}$, and $q$, representing mutant selection strength, mutation timing and cell pushing strength respectively. Median inferred parameter value is indicated by the vertical dashed line along the diagonal panels. 95% credible regions lie within the shaded region in the diagonal panels. Where no shaded region is given, this interval was the entire parameter range. **(c)** All analysed simulated sub-clonal mixing patterns (grey points) with CMFPT value of the BaseScope sub-sample (star) and posterior samples (green points). **(d)** Best-fit simulated sub-clonal pattern and parameters representing the most abundant parameter combination within the posterior distribution.

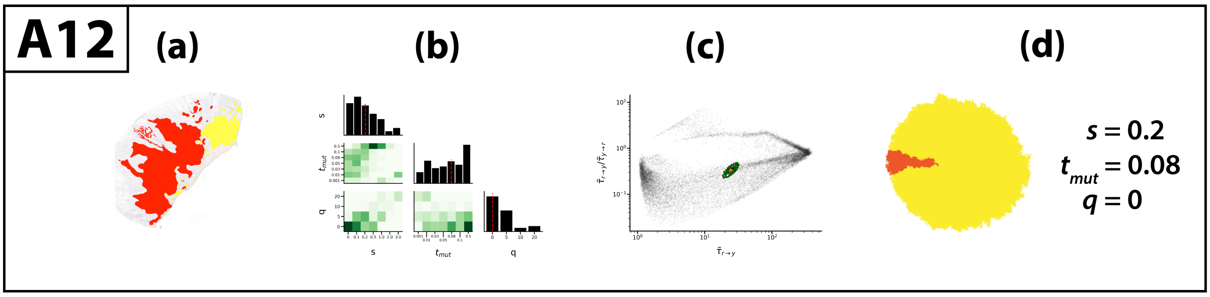

**Fig S23.** Bayesian grid-search analysis of BaseScope sample A12. **(a)** Sub-sample of sample A12. **(b)** Marginal posterior distributions of model parameters $s$, $t_{mut}$, and $q$, representing mutant selection strength, mutation timing and cell pushing strength respectively. Median inferred parameter value is indicated by the vertical dashed line along the diagonal panels. 95% credible regions lie within the shaded region in the diagonal panels. Where no shaded region is given, this interval was the entire parameter range. **(c)** All analysed simulated sub-clonal mixing patterns (grey points) with CMFPT value of the BaseScope sub-sample (star) and posterior samples (green points). **(d)** Best-fit simulated sub-clonal pattern and parameters representing the most abundant parameter combination within the posterior distribution.

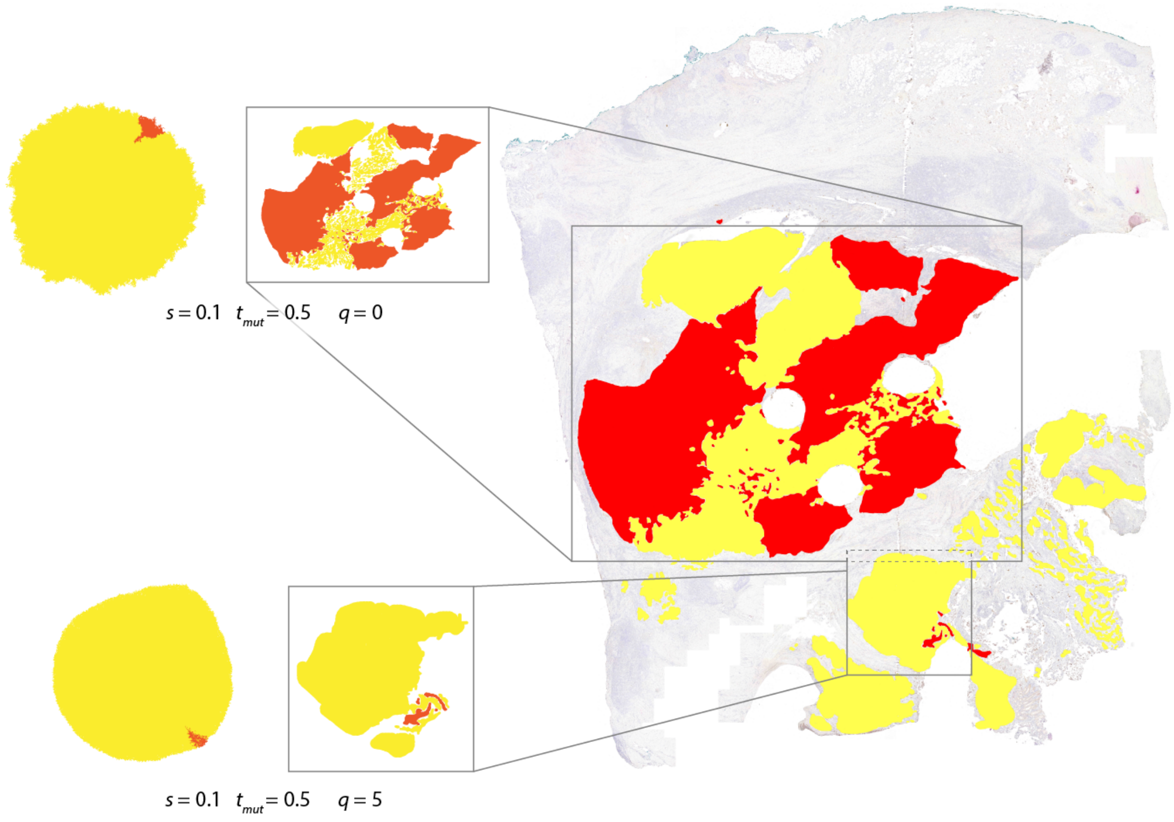

**Fig S24.** Sub-section analysis of BaseScope sample 11. Best-fit simulated sub-clonal pattern shown next to each sub-section with corresponding model parameters.

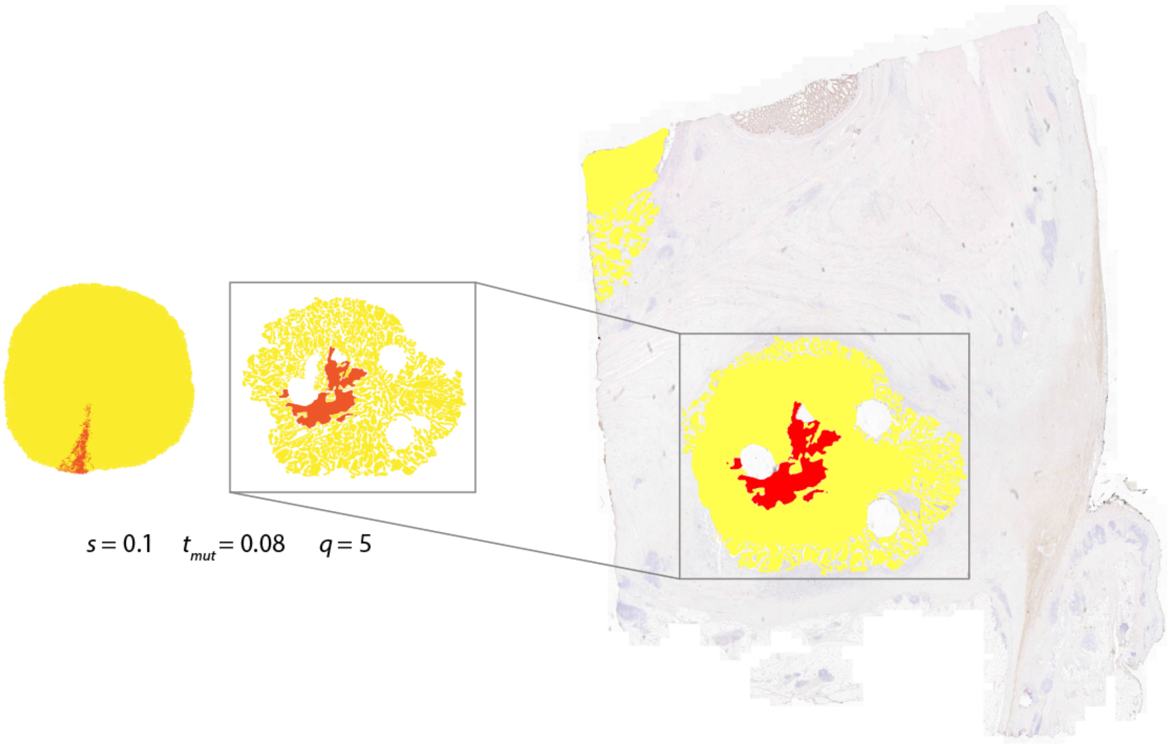

**Fig S25.** Sub-section analysis of BaseScope sample 12. Best-fit simulated sub-clonal pattern shown next to each sub-section with corresponding model parameters.

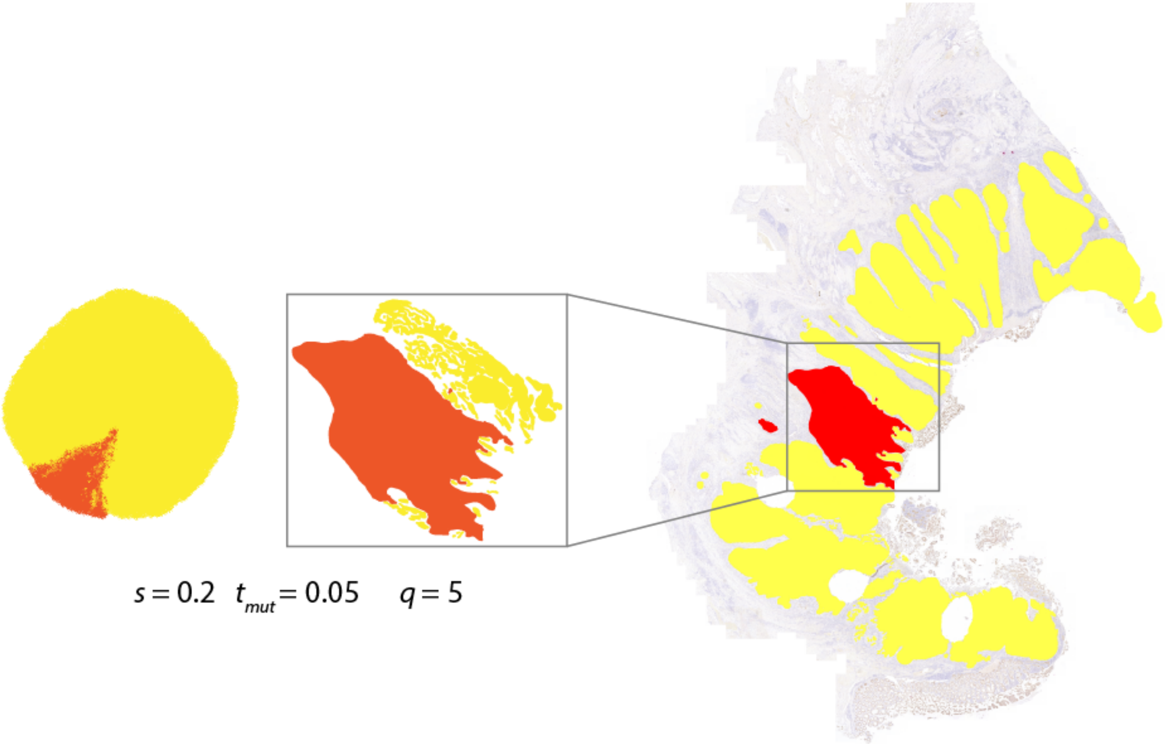

**Fig S26.** Sub-section analysis of BaseScope sample 13. Best-fit simulated sub-clonal pattern shown next to each sub-section with corresponding model parameters.

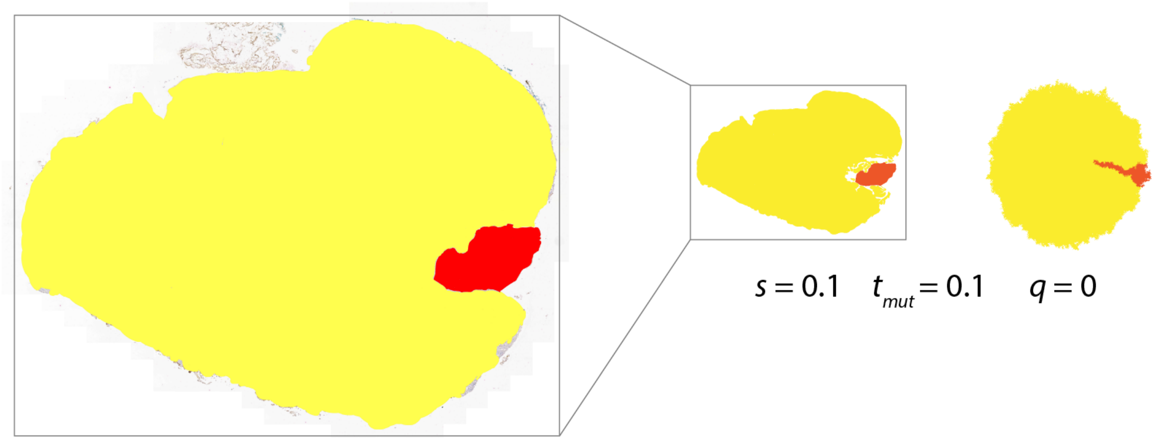

**Fig S27.** Sub-section analysis of BaseScope sample 16. Best-fit simulated sub-clonal pattern shown next to each sub-section with corresponding model parameters.

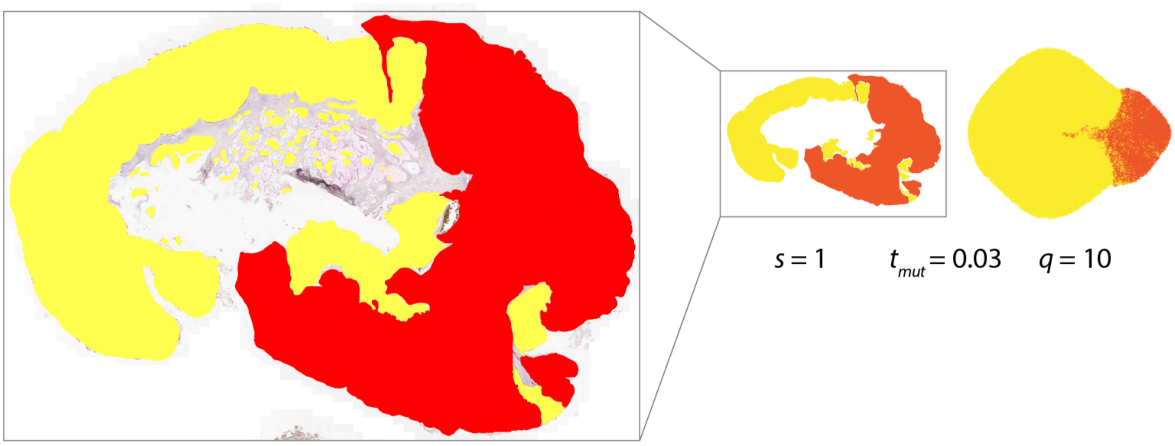

**Fig S28.** Sub-section analysis of BaseScope sample 17a1. Best-fit simulated sub-clonal pattern shown next to each sub-section with corresponding model parameters.

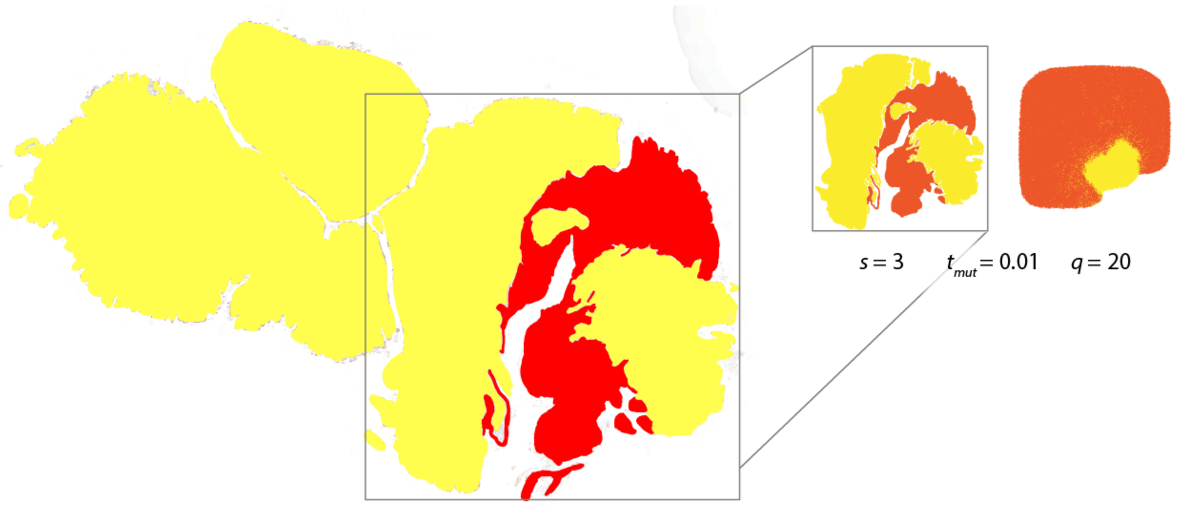

**Fig S29.** Sub-section analysis of BaseScope sample 17a2. Best-fit simulated sub-clonal pattern shown next to each sub-section with corresponding model parameters.

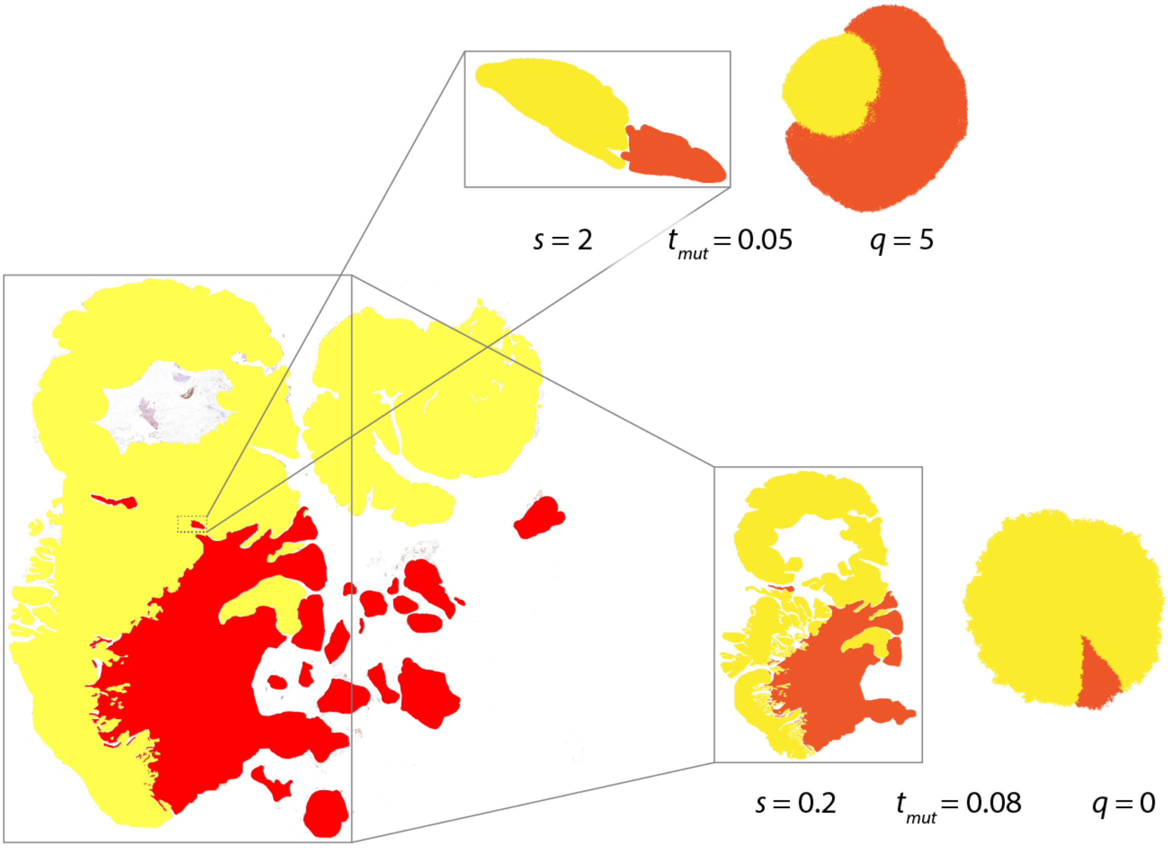

**Fig S30.** Sub-section analysis of BaseScope sample 17b. Best-fit simulated sub-clonal pattern shown next to each sub-section with corresponding model parameters.

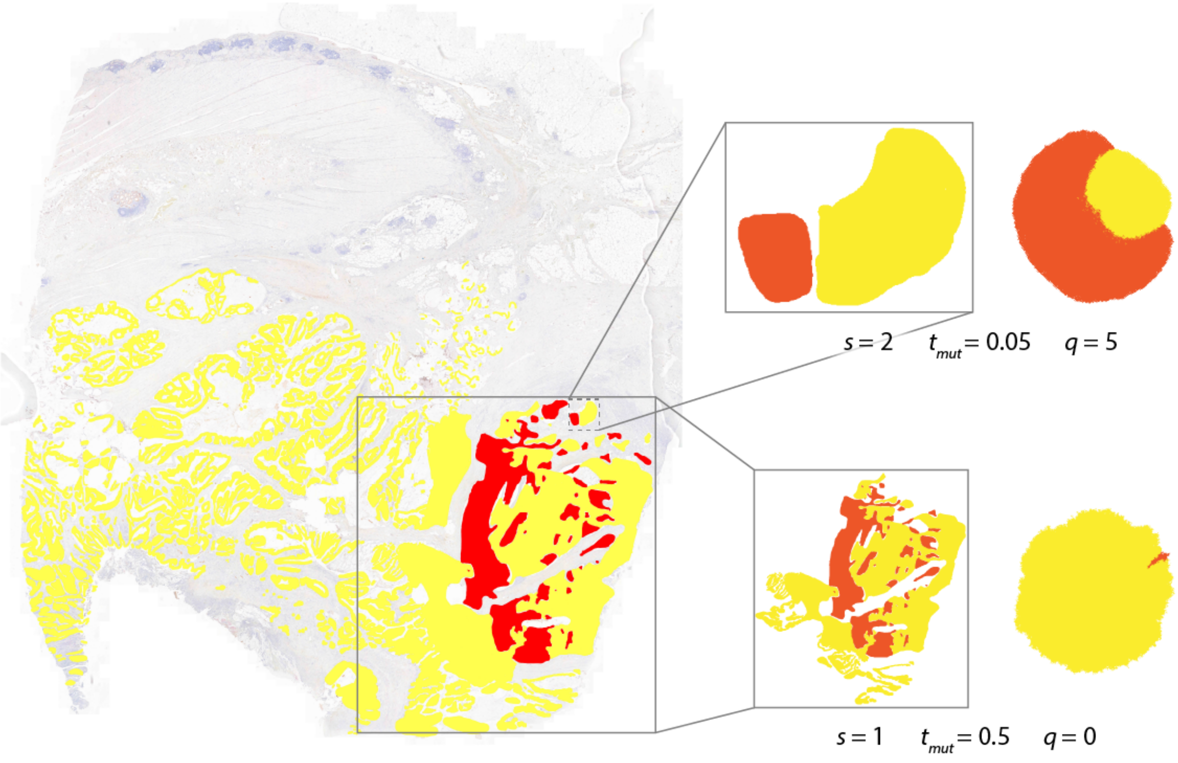

**Fig S31.** Sub-section analysis of BaseScope sample 25. Best-fit simulated sub-clonal pattern shown next to each sub-section with corresponding model parameters.

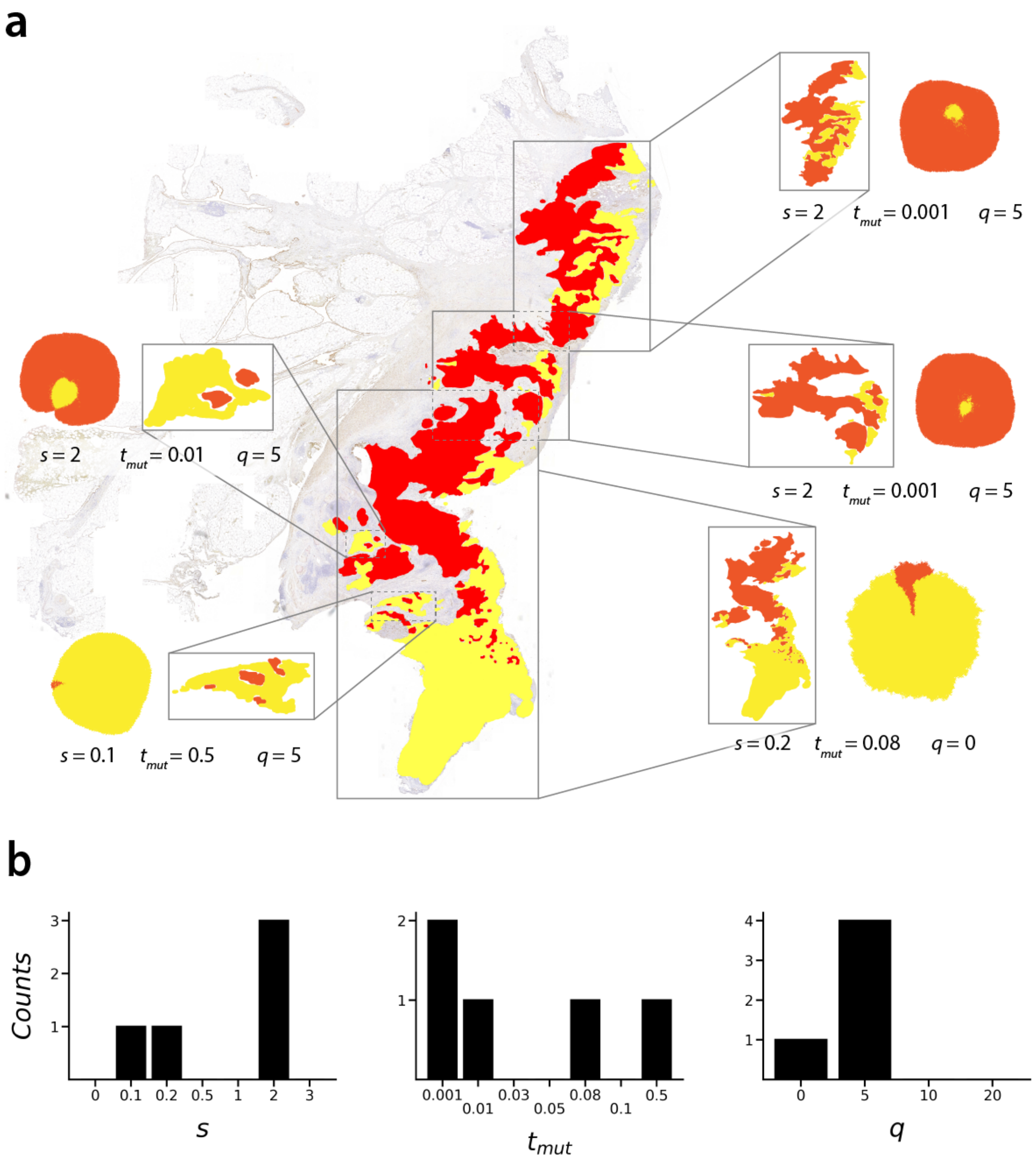

**Fig S32. Sub-section analysis of BaseScope sample 02. (a)** Sub-section analysis of BaseScope sample 02. Best-fit simulated sub-clonal pattern shown next to each sub-section with corresponding model parameters. **(b)** Marginal distributions of median inferred model parameters.

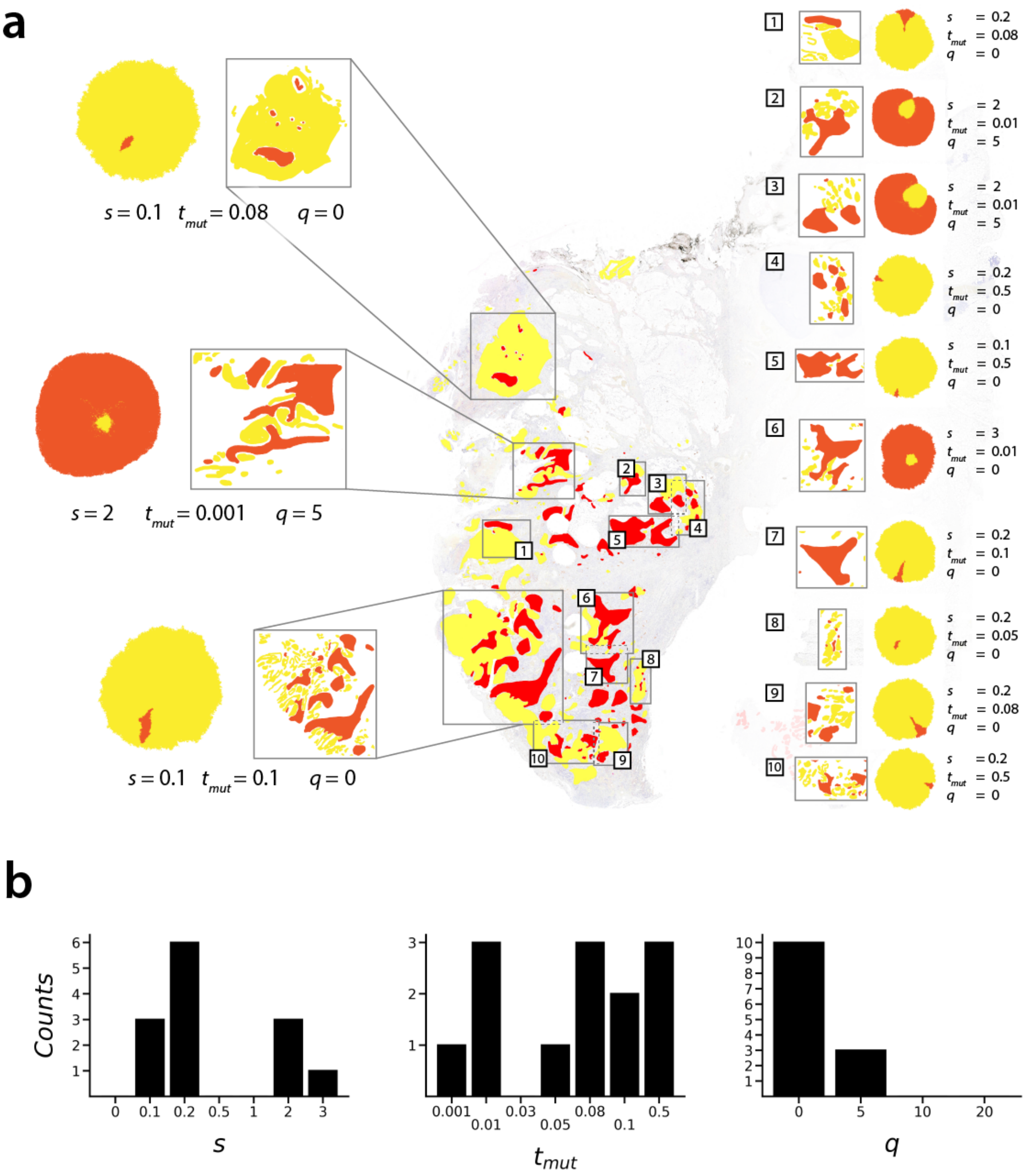

**Fig S33.** **(a)** Sub-section analysis of BaseScope sample 28. Best-fit simulated sub-clonal pattern shown next to each sub-section with corresponding model parameters. **(b)** Marginal distributions of median inferred model parameters.

| **Parameter** | **Value** | **Comments** |
|---|---|---|
| $N_{max}$ | $1 \times 10^5$ | Final number of cells |
| $\psi$ | 0.3 | Probability of cell death per birth event |
| $s$ | $\{0, 0.1, 0.2, 0.5, 1, 2, 3\}$ | Mutation selective advantage |
| $t_{mut}$ | $\{0.001, 0.01, 0.03, 0.05, 0.08, 0.1, 0.5\}$ | Fraction of total system growth before mutation |
| $q$ | $\{0, 5, 10, 20\}$ | Cell pushing chain length |
| | 200 | Min. accepted final mutant clone size |

**Table S1.** Model parameters used to generate *in silico* sub-clonal mixing patterns.

| Sample | Best-fit (3D histogram) | Best-fit (marginal median) |
|---|---|---|
| 02 (1) | (2.0, 0.001, 5) | (2.0, 0.001, 5) |
| 02 (2) | (2.0, 0.001, 0) | (2.0, 0.001, 5) |
| 02 (3) | (0.5, 0.5, 0) | (0.2, 0.08, 0) |
| 02 (4) | (3.0, 0.08, 5) | (2.0, 0.05, 5) |
| 02 (5) | (3.0, 0.01, 5) | (2.0, 0.01, 5) |
| 02 (6) | (0, 0.5, 0) | (0.1, 0.5, 5) |
| 11 (1) | (0.1, 0.5, 0) | n/a |
| 11 (2) | (0.2, 0.5, 5) | (0.1, 0.5, 5) |
| 11 (3) | (0.1, 0.08, 0) | (0.2, 0.03, 0) |
| 11 (4) | (3.0, 0.01, 20) | (0.2, 0.08, 0) |
| 12 (1) | (1.0, 0.5, 10) | (0.1, 0.08, 5) |
| 13 (1) | (1.0, 0.5, 5) | (0.2, 0.05, 5) |
| 16 (1) | (0.1, 0.5, 0) | (0.1, 0.1, 0) |
| 17a1 (1) | (0.5, 0.001, 10) | (1.0, 0.03, 10) |
| 17a2 (1) | (3.0, 0.01, 20) | n/a |
| 17b (1) | (0.5, 0.5, 0) | (0.2, 0.08, 0) |
| 17b (2) | (3.0, 0.08, 0) | (2.0, 0.05, 5) |
| 25 (1) | (3.0, 0.08, 0) | (2.0, 0.05, 5) |
| 25 (2) | (0, 0.5, 0) | (0.1, 0.5, 0) |
| 28 (1) | (0, 0.5, 0) | (0.1, 0.08, 0) |
| 28 (2) | (2.0, 0.001, 0) | (2.0, 0.001, 5) |
| 28 (3) | (2.0, 0.01, 0) | (2.0, 0.01, 5) |
| 28 (4) | (3.0, 0.01, 10) | (2.0, 0.01, 5) |
| 28 (5) | (0.2, 0.5, 0) | n/a |
| 28 (6) | (0.1, 0.5, 0) | (0.2, 0.1, 0) |
| 28 (7) | (0.2, 0.5, 0) | (0.1, 0.5, 0) |
| 28 (8) | (0.5, 0.5, 0) | (0.2, 0.08, 0) |
| 28 (9) | (2.0, 0.03, 0) | (2.0, 0.03, 0) |
| 28 (10) | (0, 0.5, 0) | (0.1, 0.1, 0) |
| 28 (11) | (3.0, 0.01, 0) | n/a |
| 28 (12) | (0.1, 0.5, 0) | (0.2, 0.1, 0) |
| 28 (13) | (2.0, 0.001, 0) | (0.2, 0.05, 0) |
| 28 (14) | (0.1, 0.5, 0) | (0.2, 0.5, 0) |
| 28 (15) | (3.0, 0.01, 20) | (0.2, 0.08, 0) |
| 34 (1) | (2.0, 0.001, 5) | (2.0, 0.001, 5) |
| 34 (2) | (3.0, 0.01, 20) | n/a |
| 34 (3) | (3.0, 0.03, 5) | (3.0, 0.03, 0) |
| 34 (4) | (0, 0.5, 0) | (0.1, 0.5, 0) |
| 34 (5) | (3.0, 0.01, 0) | (2.0, 0.001, 0) |
| 34 (6) | (3.0, 0.01, 5) | (3.0, 0.01, 0) |
| 34 (7) | (3.0, 0.03, 5) | (3.0, 0.01, 5) |
| 34 (8) | (2.0, 0.001, 0) | (2.0, 0.001, 5) |
| C537_A7 | (3.0, 0.01, 5) | (3.0, 0.01, 5) |
| C537_A10 | (3.0, 0.01, 20) | (0.2, 0.05, 0) |
| C537_A11 | (3.0, 0.03, 0) | (3.0, 0.01, 5) |
| C539_A12 (1) | (0.5,0.5,0) | (0.2,0.08,0) |

**Table S2.** Comparison of best-fit model parameters, formatted as ($s$, $t_{mut}$, $q$), as determined by the 3D histogram and marginal median methods. Sub-sample number is shown in parentheses next to BaseScope sample number. For samples whose marginal median best-fit parameters are marked as n/a, the 3D histogram method was used instead of marginal median method in the main results to determine best-fit parameters.
